# Spatiotemporal profiling defines persistence and resistance dynamics during targeted treatment of melanoma

**DOI:** 10.1101/2024.02.02.577085

**Authors:** Jill C. Rubinstein, Sergii Domanskyi, Todd B. Sheridan, Brian Sanderson, SungHee Park, Jessica Kaster, Haiyin Li, Olga Anczukow, Meenhard Herlyn, Jeffrey H. Chuang

## Abstract

Resistance of BRAF-mutant melanomas to targeted therapy arises from the ability of cells to enter a persister state, evade treatment with relative dormancy, and repopulate the tumor when reactivated. Using spatial transcriptomics in patient derived xenograft models, we capture clonal lineage evolution during treatment, finding the persister state to show increased oxidative phosphorylation, decreased proliferation, and increased invasive capacity, with central-to-peripheral gradients. Phylogenetic tracing identifies intrinsic- and acquired-resistance mechanisms (e.g. dual specific phosphatases, Reticulon-4, CDK2) and suggests specific temporal windows of potential therapeutic efficacy. Using deep learning to analyze histopathological slides, we find morphological features of specific cell states, demonstrating that juxtaposition of transcriptomics and histology data enables identification of phenotypically-distinct populations using imaging data alone. In summary, we define state change and lineage selection during melanoma treatment with spatiotemporal resolution, elucidating how choice and timing of therapeutic agents will impact the ability to eradicate resistant clones.

**Statement of Significance:** Tumor evolution is accelerated by application of anti-cancer therapy, resulting in clonal expansions leading to dormancy and subsequently resistance, but the dynamics of this process are incompletely understood. Tracking clonal progression during treatment, we identify conserved, global transcriptional changes and local clone-clone and spatial patterns underlying the emergence of resistance.

---

Tumors are collections of heterogeneous clonal populations that continually evolve under selective pressures imposed upon them by the necessity of vying for survival in a resource constrained environment. Subjecting the tumor ecosystem to targeted treatment introduces a bottleneck, either selecting for those populations with pre-existing resistance characteristics, or driving the evolution of novel resistance mechanisms. BRAF-mutant melanoma often follows a pattern of initial dramatic response to targeted treatment, leaving minimal residual or no detectable disease, followed at varying intervals by regrowth of resistant tumor.^1–3^ Persister cells have been defined as small, transient, drug-tolerant populations that maintain tumor viability until larger, drug-resistant populations regrow.^4^

The persister state has been observed in melanoma, presenting an evolutionary bottleneck through which tumors inevitably progress in their clinical trajectory toward treatment failure.^5^ The temporal dynamics and specific pathways leading into and out of the persister state remain unclear. Due to challenges of longitudinal sample acquisition, much of what is known of this state comes from *in vitro* studies, which have identified reproducible characteristics including increased expression of histone demethylases, associated chromatin remodeling, and slow or absent proliferation, as well as alterations in cellular respiration.^4,6,7^ Recent evidence from single cell sequencing studies has highlighted heterogeneity *within* persister populations, with only rare sub-populations possessing the ability to proliferate under the pressure of therapy.^8^ Further evidence of the importance of clonal heterogeneity has been shown in patient derived xenograft (PDX) studies with single-cell sequencing and multiplexed immunohistochemistry to reveal multiple, spatially-distinct, drug-tolerant states.^9^ Indeed, PDX models allow tumor assessment across time, providing a potent approach to interrogate resistance dynamics. However, heterogeneity is influenced by spatial localization, and the *in situ* dynamics of clonal heterogeneity during treatment remain poorly understood. Elucidating these processes requires not only an *in vivo* model and longitudinal treatment design, but the use of techniques capable of resolving clonal identities with high spatial resolution.

To address this gap, here we characterize the *in vivo* transcriptional and evolutionary trajectory of *BRAF*-mutant melanomas using PDX models treated with the combination targeted agents dabrafenib and trametinib. Mice were treated through minimal residual disease (MRD) and subsequent tumor regrowth with sampling of the tumors at multiple time points during the treatment course. This was repeated in two separate PDX models, deriving from different patients but following parallel treatment response trajectories. We employ a spatial transcriptomics (ST) assay and deep learning techniques on associated hematoxylin & eosin (H&E)-stained slides to capture gene expression- and phenotypic-profiles that reflect disease state, while maintaining intact tissue structure. We also introduce a comprehensive computational pipeline for processing and integration of these joint spatiotemporal data. This combination of dense time point sampling with spatially resolved profiling provides unusual access to the dynamics underlying tumor evolution during treatment, showing that: all cells have the potential to enter the persister state; there are conserved spatial patterns within treated tumors including clonal neighbor preferences and central-to-peripheral gradients; resistance is due to a combination of intrinsic- and acquired-mechanisms; there are stereotypic temporal changes in cell state during treatment; and deep learning on H&E-stained images can help track these cell states even in the absence of molecular data.

## Results

### Spatial profiling quantifies tumor size more accurately than in situ measurement

Investigation of the minimal residual disease state requires accurate measurement of tumor size during progression from pre-treatment, during shrinkage to minimal measured volume, and during subsequent regrowth. Traditional methods rely on *in vivo* tumor volume assessment using calipers, with size reported as (width^2^ x length)/2. However, such estimates of gross volume do not distinguish viable tumor relative to mouse stroma and necrotic material. Comparison of *in vivo* tumor volume measurements to estimates from whole tumor resected spatial transcriptomics sections demonstrates that gross measurement can misidentify landmark moments in the treatment course– specifically the maximally shrunken timepoint, which has been presumed dominated by resistant clones in a persister state. Resection-based spatial analysis allows for filtering of mouse stroma and therefore more precise tracking of tumor size over time, improving association of expression signatures with moments before, during, and after the evolutionary bottleneck imposed by treatment (Figure 1a,b).

**Figure 1.**
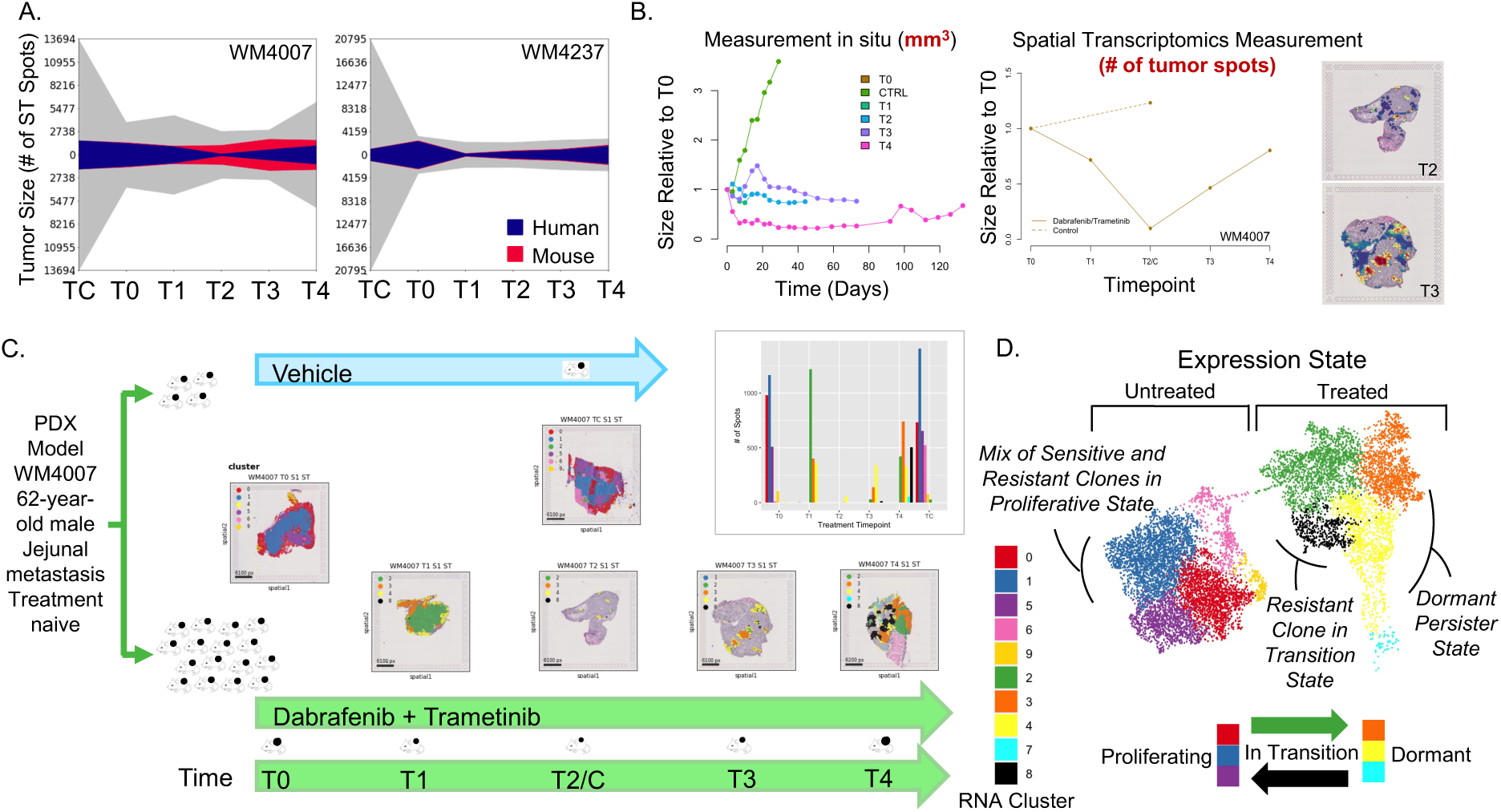
Spatial transcriptomics allows accurate measurement of tumor size and captures evolution of RNA-based clusters across treatment. A. Tumor size for assayed samples as captured in situ by calipers vs. whole resection ST measurement for both models WM4007 and WM4237. Blue area represents the number of ST spots predominantly comprised of human tissue (tumor). Red area represents ST spots for surrounding mouse tissue. Grey area represents size as captured by calipers, estimated from maximum tumor diameter. B. (left) WM4007 tumor growth curves as measured in situ versus (center) size at resection estimated by viable tumor in the ST image. (right) H&E-stained sections of time points 2 and 3, colored by the prevalence of human reads, illustrating that the minimal size is at time point T2, not T3 as would be suggested by in situ measurement. C. Summary of ST sections for WM4007 at each time point, colored by RNA-derived clusters. Bar chart shows the proportions of RNA clusters at each time point. D. UMAP projection of RNA clusters of WM4007 showing transition from a proliferating (left) into a dormant (right) expression state, with eventual re-entry into proliferative state by the resistant lineage (black). Abbreviations: Spatial Transcriptomics (ST)

We perform longitudinal treatment studies on two models: WM4007 (62-year-old man, jejunal metastasis) and WM4237 (29-year-old woman, brain metastasis) with samples taken prior to treatment (T0), at four time points during the treatment course (T1-4), and from an untreated control mouse harvested concurrently with T2 (TC). In model WM4007, gross measurements imply that at T3 the tumor remains in a maximally shrunken, quiescent state. However, ST measurements reveal that at T3 the ratio of tumor to stroma is already increasing and that the maximally shrunken sample is in fact at T2. It follows that at T1 the tumor is still shrinking and therefore this sample gives unambiguous information about dynamics into the quiescent, persister state.

### Evolution during treatment exhibits stereotyped progression in expression and spatial organization

Although each tumor sample grows on the back of an individual mouse, for each model we observe a noteworthy continuity in behaviors across samples. There are consistent therapy-induced shifts in expression across samples, as observed from expression-based clustering of the pool of ST spots from all samples of a given model (see Methods). In both WM4007 and WM4237, the pre-treatment time points consist of 5 distinct expression clusters, but initiation of treatment causes sufficient changes that the T1 samples are each composed entirely of 4 new clusters. These new clusters continue to be observed at later time points and comprise the dominant expression states during minimal residual disease. Such stereotyped behavior demonstrates the low stochasticity in treatment-induced alterations in global gene expression during these evolutionary stages (Figure 1c, Supplementary Figure 1). Viewed on the UMAP projection, RNA clusters in the untreated samples (left) segregate from those that have been treated (right). In WM4007, there is a transition cluster (2, green) at the bridge between proliferating and dormant cells demonstrating passage into dormancy, and a similar cluster (8, black) corresponding to re-emergence of a proliferating, resistant population (Figure 1d), as discussed further below.

Differentially expressed genes (DEGs) between pre-treatment (T0) and early-treatment (T1) samples show common treatment-induced changes shared across the two models (Supplementary Table 1). Enrichment analysis of these DEGs reveals highly significant alterations in cell cycle, oxidative phosphorylation, and ribosomal pathways upon initiation of treatment (p<.00001 for all), with a return toward baseline state in later time points (Figure 2a, Supplementary Figure 2). Recurrent patterns are also apparent in the spatial organization of RNA-derived clusters, with specific expression states tending to occupy central tumor regions and others at the periphery or tumor-stromal boundary. This is true of both pre- and post-treatment samples as shown in Figure 2b. The mean distance to tumor boundary differs significantly between the blue and the red clusters (Clusters 0 and 1) in T0 (p<.00001). In T1, the mean distance to tumor boundary differs significantly between the green and orange clusters (2 and 3, p<.00001). Individual genes and gene sets also exhibit significant correlation with distance from the tumor boundary (Figures 2c,d, see Supplementary Table 2 for complete list of correlations by time point with p-values). For example, mitochondrial gene expression is negatively correlated with distance to the nearest tumor edge (greater expression on periphery), while mitophagy and structural maintenance of chromosomes (SMC) gene sets show greater expression more centrally. These behaviors are observed regardless of whether the sample is treated. In contrast, for other gene pathways the spatial gradient reverses direction upon the initiation of treatment. Glycolysis as well as interferon stimulated genes (ISG) are more peripherally active pre-treatment, but early treatment samples show a drastic increase in central expression. Conversely, oxidative phosphorylation, lysosome, ribosome, and invasive pathways move from a central to peripheral pattern of activation.

**Figure 2.**
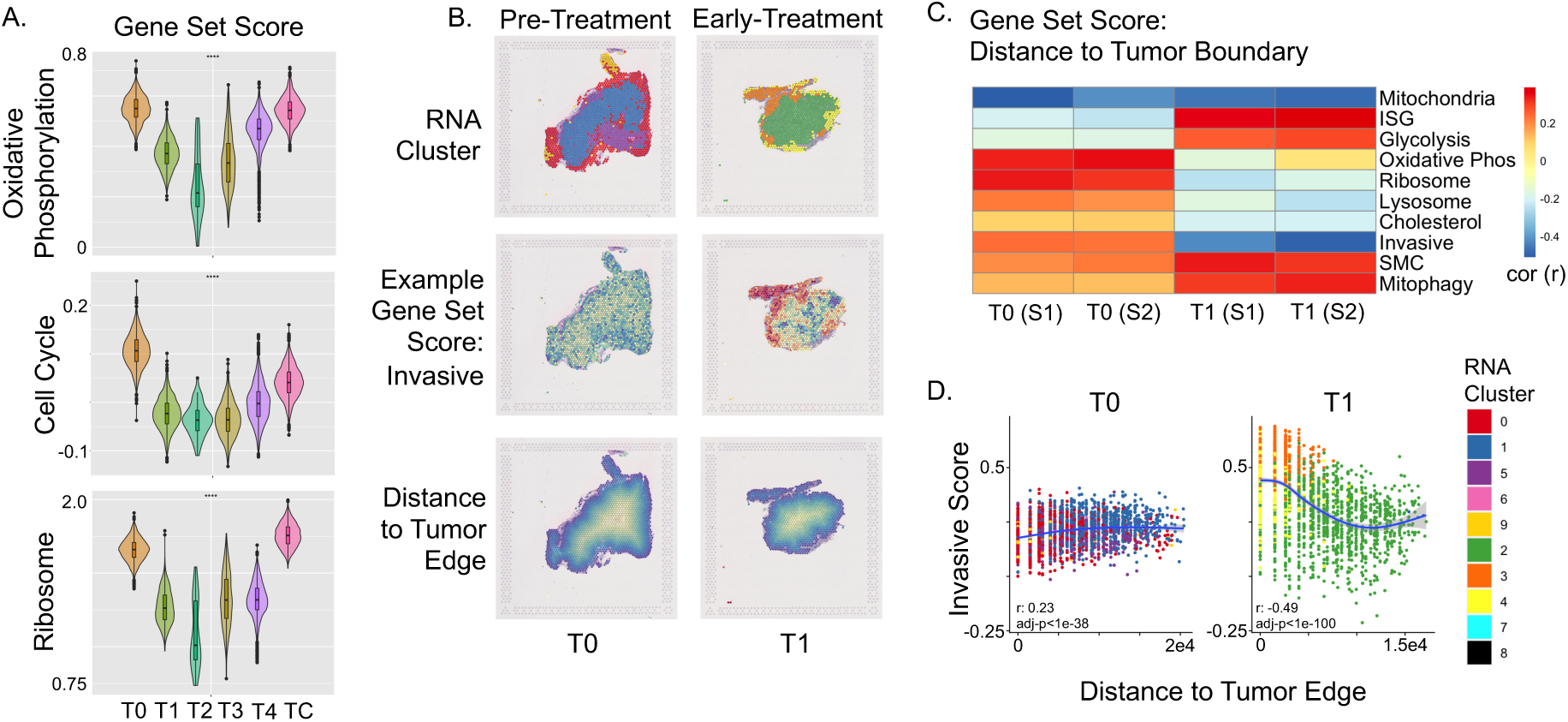
Evolution during treatment follows stereotyped progression in expression and spatial organization. A. Pathway enrichment analysis of DEGs before and after the initiation of treatment (WM4007) demonstrates initial significant down-regulation of the cell cycle, oxidative phosphorylation, and ribosome pathways with return toward baseline in the late time points. B. Recurrent spatial patterns of RNA-derived clusters show specific expression states tending to occupy central tumor regions and others at the periphery or tumor-stromal boundary, both in pre- and post-treatment samples. C. Correlation of expression of gene sets with distance from the tumor boundary, as a function of treatment timepoint. Replicate tumor slices are indicated by S1 and S2. For example, glycolysis activity is highest in the tumor periphery pre-treatment, but reverses to be higher in the tumor core upon initiation of treatment. D. The gene set score reflecting tumor invasive capacity demonstrates a positive correlation with distance from tumor edge in the pre-treatment samples but reverses with treatment. Dots indicate individual ST spots. Colors distinguish spots by their RNA expression clusters, as in Figure 1. Abbreviations: Differentially Expressed Gene (DEG)

### Spatial distribution of clusters identifies a recurrent boundary population with gradients in cell cycle, mitochondrial, lysosomal, and ribosomal expression

Recurrent spatial patterns can be observed between cells in different expression states. For example, in several sections among the treated specimens, RNA-defined clusters form concentric rings with ordering conserved along the spatial nesting such that an interior orange cluster (4) is surrounded by yellow (7) which is surrounded by cyan (3) (Figure 3a, best seen in T2 and T4). The UMAP projection demonstrates that clusters that neighbor each other spatially also neighbor each other in expression space. For example, Figure 3a,b illustrates this trend, with the olive green (2), blue (1), and pink (5) clusters occupying the central areas of the untreated sample (Figure 3a, top left) and also bordering each other in the UMAP (Figure 3b, center), while the red (0) and purple (6) cluster together in the periphery of the tumor as well as on the UMAP projection.

**Figure 3.**
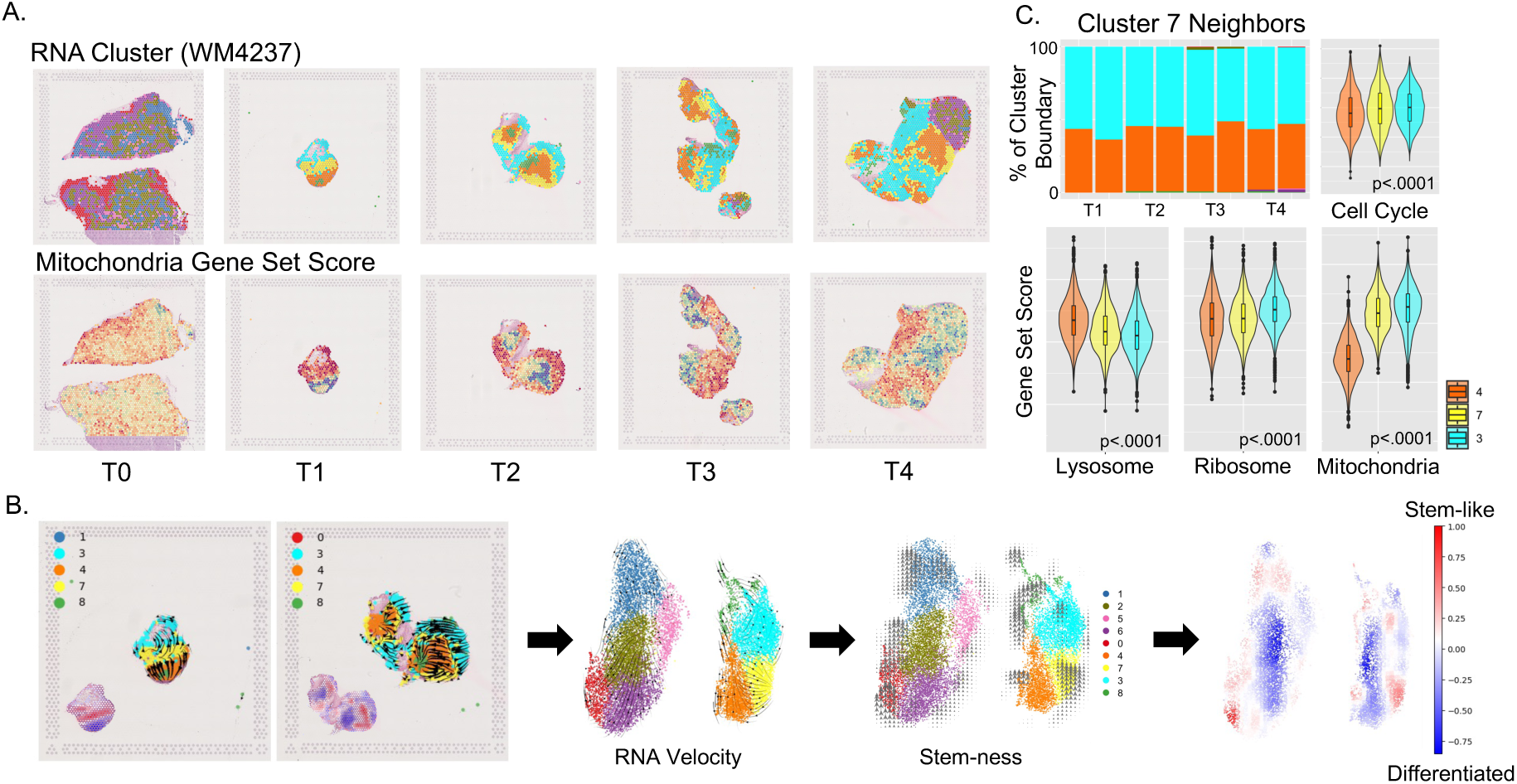
Spatial distribution of clusters identifies a recurrent boundary population. A. Spatial layout of RNA-defined clusters across timepoints for WM4237. Nearly concentric structures are observable, particularly in T2 and T4, with cluster ordering conserved along the spatial nesting: orange – yellow – cyan from internal to external. The mitochondrial gene set score follows the spatial gradient, reflecting patterns in the mechanism of cellular respiration. B. (Left) Spatial layout of RNA-defined clusters for time points 1 and 2, with arrows representing RNA velocity. Arrows point from more stem-like toward more differentiated clusters. inlay with stem-ness colored red. (Left center) UMAP projection of RNA-defined clusters. Overlying arrows show RNA velocity. (Right center) Demarcation of stem-like regions, as inferred from RNA velocity. (Right) UMAP projection of stemness vs. differentiation. C. Frequencies of expression clusters that neighbor the most stem-like cluster (7, yellow). It is nearly universally surrounded by clusters 3 and 4 (cyan and orange) throughout timepoints T1-T4. Violin plots demonstrate gradients in expression along the orange to yellow to cyan direction (p-values by Kruskal-Wallis test). Abbreviations: Uniform Manifold Approximation and Projection (UMAP)

More generally, we can quantify recurrent patterns where one cluster is consistently interposed between two specific neighbor clusters and the “sandwiching” clusters rarely co-occur without the third cluster in between. For example, in model WM4237 quantification of neighbor-neighbor frequencies identified a consistent boundary cluster (yellow, 7) separating the orange cluster (4) and cyan cluster (3) (Figure 3c). As described above, the orange cluster tends to occur centrally within the tumor while the cyan cluster tends to occur peripherally. The enrichment of boundaries between cluster 7 and clusters 3 and 4 is statistically significant (Mann-Whitney U test, p<.00001). Differentially expressed genes between this boundary cluster and its neighbors are listed in Supplementary Table 3. Enrichment analysis on this gene set demonstrates gradients across the boundary in several processes, including oxidative phosphorylation (p<.0001), mitochondrial ATP synthesis (p<.001), and ribosomal pathways (p<.0001). Comparison of gene set scores between the boundary cluster and its neighbors confirmed these expression gradients, providing further evidence to support the spatial patterns identified based on distance-to-tumor edge (ex. lysosome, ribosome, mitochondria, p<.0001 for all) (Figure 3c).

RNA velocity measurements derived from the ratio of spliced to unspliced mRNAs demonstrate spatial gradients in states related to cell differentiation.^10^ RNA velocities indicate that the yellow boundary cluster (7) is the least differentiated of these three clusters (Figure 3b). Its centrally-prevalent neighbor, the orange cluster (4), is shifted toward a quiescent persister state, as evidenced by decreased cell cycle, mitochondrial, and ribosomal gene set scores. Of note, the lysosome pathway is upregulated in the more quiescent populations. In contrast, differentiation of the yellow (7) cluster toward its peripherally-prevalent neighbor, the cyan cluster (3), yields opposite trends in each of these pathways. This suggests that the transition from the persister to the proliferating, resistant state may be triggered by signals from the tumor periphery.

### Metabolic plasticity, decreased proliferation, and increased invasive capacity mark the transition into the persister state

Temporally-resolved transcriptional profiling identifies global shifts from pre-treatment proliferation to quiescent persistence at minimal residual disease, to the emergence of proliferation in resistant cell populations. To further clarify relationships among cell states that occur in different regions at the same time point, we utilize pseudotemporal ordering, based on the comparison to a gene set reflecting invasive capacity (see Methods) as a proxy for progression of transcriptional change.

Comparing the temporal and pseudotemporal progression of clusters illustrates the heterogeneity of individual pathways during treatment. In Figure 4a, spot expression values for three pathways are plotted vs. pseudotime, with spots color-coded by RNA cluster. Gene set scores reflecting spot-level tumor characteristics are plotted on the y-axes. Time point T1 in model WM4007 is the moment after treatment starts when the tumor is shrinking toward the persister state. There is a central-peripheral spatial gradient in several pathways that reverses from T0 to T1 (Figure 2c). This spatial gradient matches the pseudotemporal progression from cluster 2 (green, transition cluster in Figure 2d) toward cluster 4 then cluster 3 (yellow and orange) as can be seen in Figure 4a,b. The pseudotemporal and spatial progression is characterized by a shift toward aerobic respiration, as evidenced by consistent decreases in the ratio of glycolysis to oxidative phosphorylation, increases in invasive capacity, and slowing of the cell cycle (Figure 4a). Cell cycle decreases upon treatment and increases at later times, whereas invasive capacity follows the opposite trend (Figure 4c). The sparse population of T2 is dominated by cluster 4 (yellow) but T3 exhibits a pseudotemporal transition back toward RNA cluster 3 (orange) (Figure 4a). In T4, the ratio of glycolysis to oxidative phosphorylation has returned to pre-treatment levels. Invasiveness remains elevated above its baseline in clusters 2 and 3, but is reduced in the newly emerged RNA cluster 8 (black). Intriguingly, this resistant cluster 8 has lost the increased invasive capacity of other clusters, but has acquired an increased proliferative rate as reflected in the cell cycle score. These trends in cell cycle and invasion are consistent across both models (Figure 4c). However, a caveat is that by T1 in WM4237, transition to the persister state has already occurred. Therefore, there is no observable cluster corresponding to the green transition cluster 2 of WM4007 (Supplementary Figure 3).

**Figure 4.**
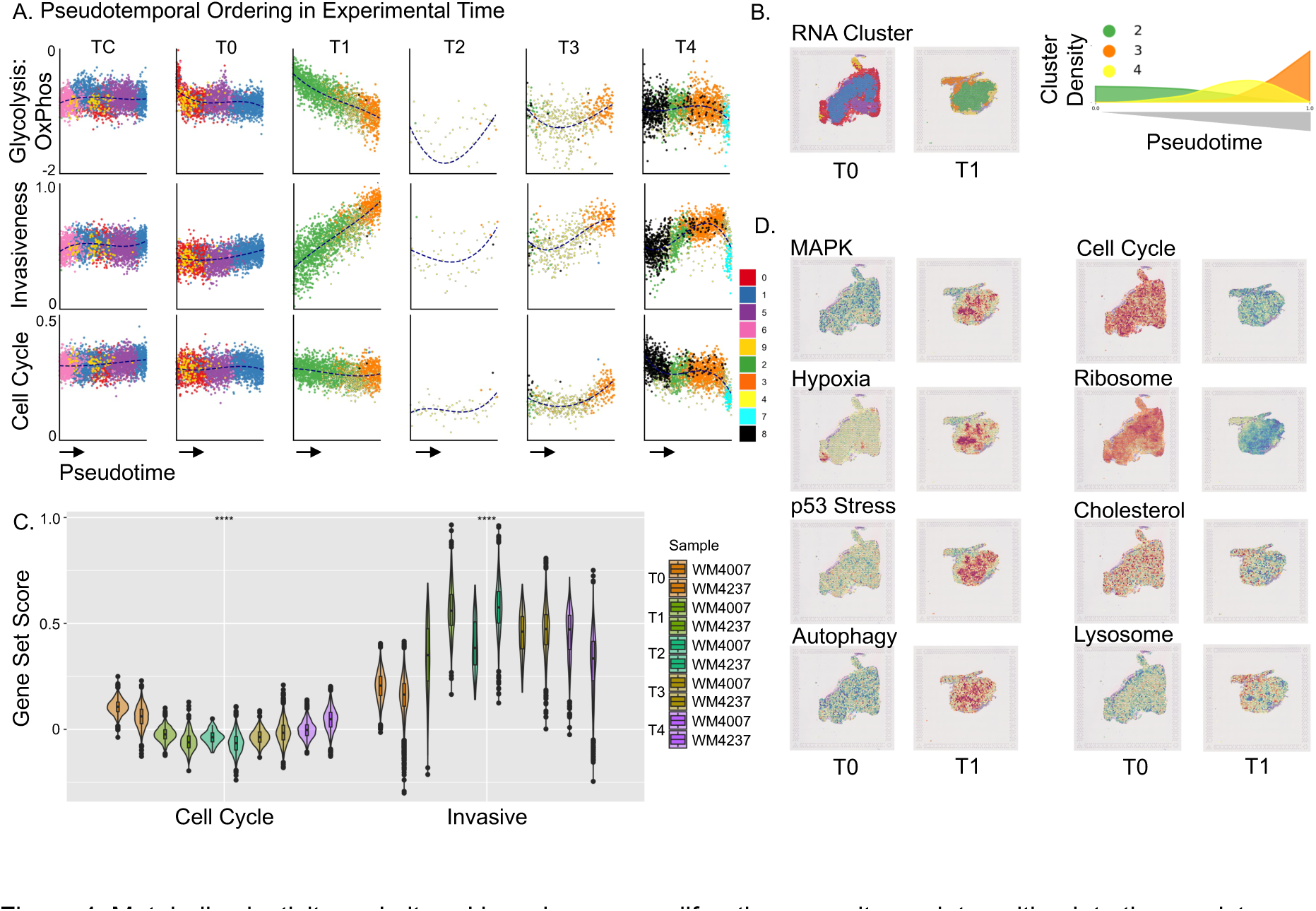
Metabolic plasticity and altered invasiveness-proliferation capacity mark transition into the persister state. A. Expression of gene sets by pseudotemporal ordering to reflect transcriptional change, graphed for each timepoint. In each graph, points show ST spots vs. pseudotime, which is defined using invasive capacity as a reference (see Methods). Gene sets: (Top) glycolysis/oxidative phosphorylation (log), (Middle) invasive capacity, and (Bottom) cell cycle. Colors indicate RNA-defined clusters. Dashed lines indicate running averages. Treatment induces a shift from glycolysis toward oxidative phosphorylation as pseudotime progresses from cluster 2 to 4 to 3 (green to yellow to orange). B. (left) Spatial layouts of RNA-derived clusters before and after the initiation of treatment demonstrate concentric patterns. (right) Cluster densities after treatment, as a function of pseudotime. C. Cell cycle and invasive capacity expression in models WM4007 and WM4237 as a function of timepoint. D. Spatial layout of additional gene pathways in WM4007, showing altered expression both in response to treatment (T0 vs. T1) and along the central-to-peripheral axis.

Several additional pathways demonstrate altered expression in the transition cluster captured in WM4007 T1. There is strong up-regulation of MAPK, hypoxia, p53 stress, and autophagy (Figure 4d). In contrast, the cholesterol and lysosome pathways demonstrate suppression in the transition cluster 2 and upregulation in the peripheral clusters (3 and 4). Cell cycle and ribosome show strong global down-regulation upon initiating treatment, though with some rebound in ribosome in cluster 3. These results are in line with prior studies regarding key pathways in melanoma progression and therapeutic resistance, adding previously inaccessible temporal and spatial information. For example, upregulation of the autophagy-lysosome axis is a known response to kinase inhibition, including in BRAF-mutant melanoma treated with targeted inhibition.^11^ The addition of specific inhibition of autophagy as part of combination therapy has been proposed, with in vitro studies showing promise.^12–14^ Similarly, cholesterol and ribosome biosynthesis pathways have been linked to melanoma plasticity and metastatic potential.^15,16^ These spatiotemporal variations suggest that success in a combined targeted therapy approach will benefit from precise elucidation of when and where these pathways are most active.

### Phylogeny tracking reveals global transcriptional state change followed by lineage-specific survival through the treatment bottleneck

Copy number variants (CNVs) accumulate in the genome during the course of cell proliferation, providing signatures of phylogenetic lineage. By generating CNV profiles using inferCNV^17^ on spot transcriptome data, we are able to track clonal phylogenies with spatial and temporal resolution. Cell states within these phylogenies can be evaluated simultaneously from each spot’s transcriptome. Clustering by CNV profiles resulted in the identification of 10 distinct clonal lineages in WM4007 and 12 in WM4237. Figure 5a shows the WM4007 RNA-derived UMAP projection, colored by RNA cluster, CNV lineage, resistance potential, and time point, providing an overview of the relationships between transcriptional state and phylogenetic lineage. Figure 5b re-displays these same data to show time on the x-axis and CNV lineages on the y-axis, arranged from treatment-sensitive at the top to treatment-resistant at the bottom. Coloring of the spots by RNA cluster highlights the transcriptional changes occurring over time in each lineage. In both models, all phylogenetic lineages existing in the pre-treatment time point survive the initiation of treatment, though lineages vary in population size at T1. This analysis demonstrates that the ability to enter the persister state is not limited to a small subpopulation of cancer cells, as has been assumed previously.^4,18^ Rather, there is a global shift where all lineages transition into the persister state characteristic of minimal residual disease (Supplementary Figure 4).

**Figure 5.**
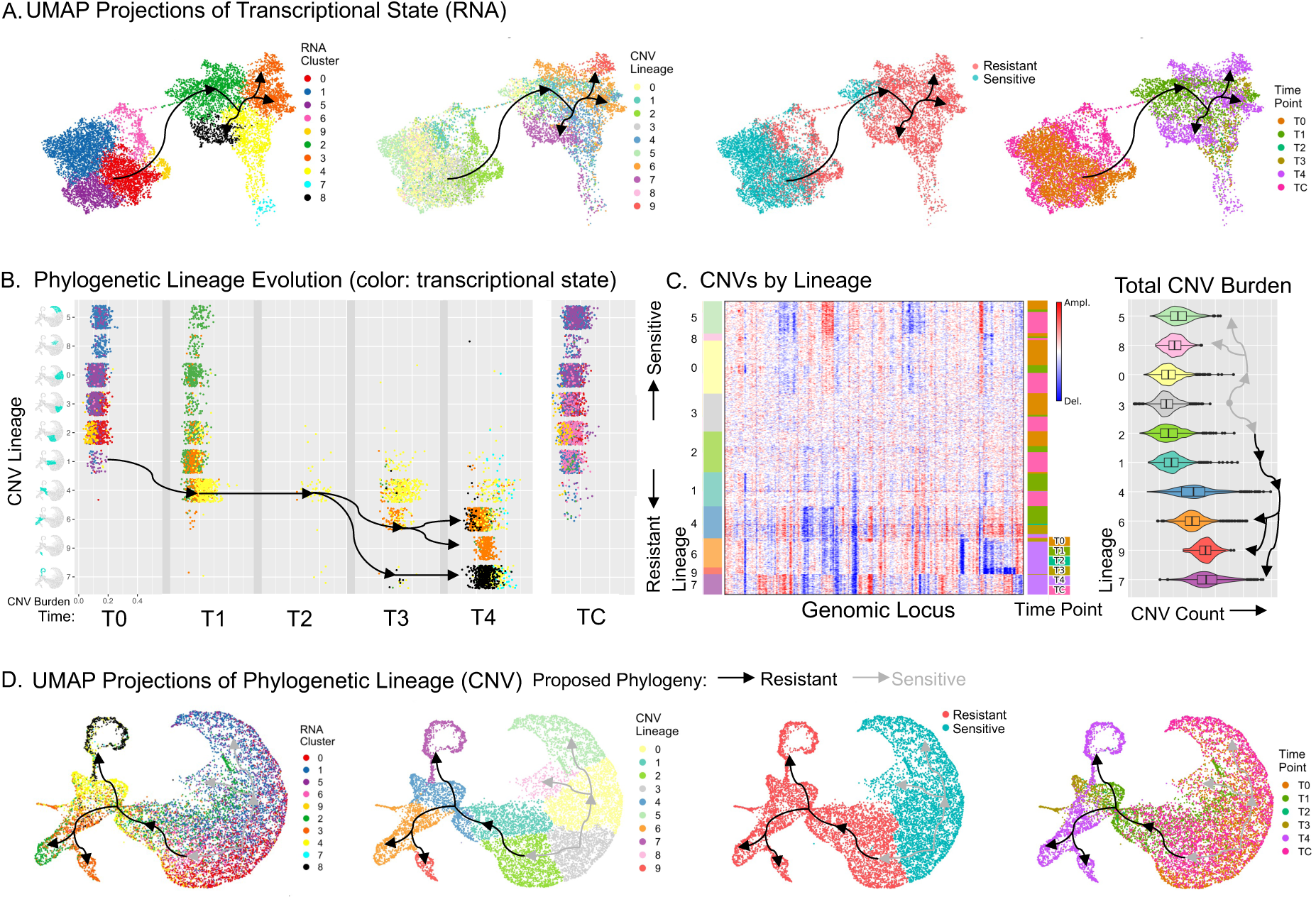
Phylogeny tracking reveals global transcriptional state change followed by lineage-specific survival through the treatment bottleneck (WM4007). A. RNA-derived UMAP projection, colored by RNA cluster, CNV lineage, resistance potential, and time point (left to right). B. Phylogeny of CNV lineage evolution from pre-treatment to the re-emergent, resistant state. Each dot corresponds to an ST spot. Color indicates RNA cluster, showing transcriptional state changes for each lineage during treatment. C. (Left) Heatmap showing CNVs for each lineage, plotted for each spot and timepoint. (Right) Total CNV burden for each lineage across all spots, reflecting direction of evolution. D. CNV-derived UMAP projection, colored by RNA cluster, CNV lineage, resistance potential, and time point (left to right), demonstrating lineage bottleneck with emergence into two divergent branches by time point 4. Abbreviations: Uniform Manifold Approximation and Projection (UMAP), Copy Number Variant (CNV)

After this global transcriptional state change, the ability to survive through the persister state and re-emerge into a proliferating clone varies by lineage. The phylogenetic distance between lineages is reflected in the heatmap and UMAP projections of CNV profiles (Figures 5c,d). A group of related lineages (CNV 0, 3, 5, 8) occupy the right half of the CNV UMAP and demonstrate behavior expected of treatment sensitive clones– they are present in T0 but extinguished by T2 (Figure 5d left-middle panel). At T1 all of these lineages exhibit transitional expression patterns (green RNA cluster), and some spots from these lineages have reached the persister state (orange and yellow RNA clusters). Cosine distances calculated between CNV lineages demonstrate that the treatment sensitive clones lack a strong connection with any surviving lineage, indicating they have been extinguished by treatment (Supplementary Figure 5, left).

Intriguingly, the majority of CNV lineage 1 has entered the persister state by T1, and a small population from this lineage survives the treatment bottleneck and begins regrowth by T4, suggesting this lineage harbors intrinsic therapeutic resistance. The CNV UMAP projection shows a close relationship between lineages 1 and 4, the predominant population surviving through the treatment bottleneck. Additionally, lineage 4 carries a greater CNV burden than lineage 1. This begs the question of whether the distinguishing variants drive novel (acquired) resistance mechanisms or whether resistance is due to the same mechanisms as in lineage 1 (see section on resistance below, Figure 6a). Lineage 4 is the predominant tumor population at MRD time point T2. This evolutionary bottleneck can be observed in the narrowing in the CNV UMAP as the tumor evolves from right to left. In line with prior studies, we find that the regrowing tumors maintain clonal complexity^19^ and indeed accrue further heterogeneity as can be seen in phylogenetic bifurcations at T3 and T4. New CNV lineages 6 and 7 are detectable at these times. These exhibit transcriptional changes from the MRD state, with 6 transitioning from the yellow to the previously seen orange RNA cluster, while lineage 7 begins to repopulate the tumor in a novel transcriptional state (black, RNA cluster 8). Finally, in T4, lineage 6 spawns lineage 9 and itself begins to transition to the novel black transcriptional state. This proposed resistance phylogeny is illustrated by arrows overlying the projections of the data in the UMAPs (Figure 5d, left-middle panel).

**Figure 6.**
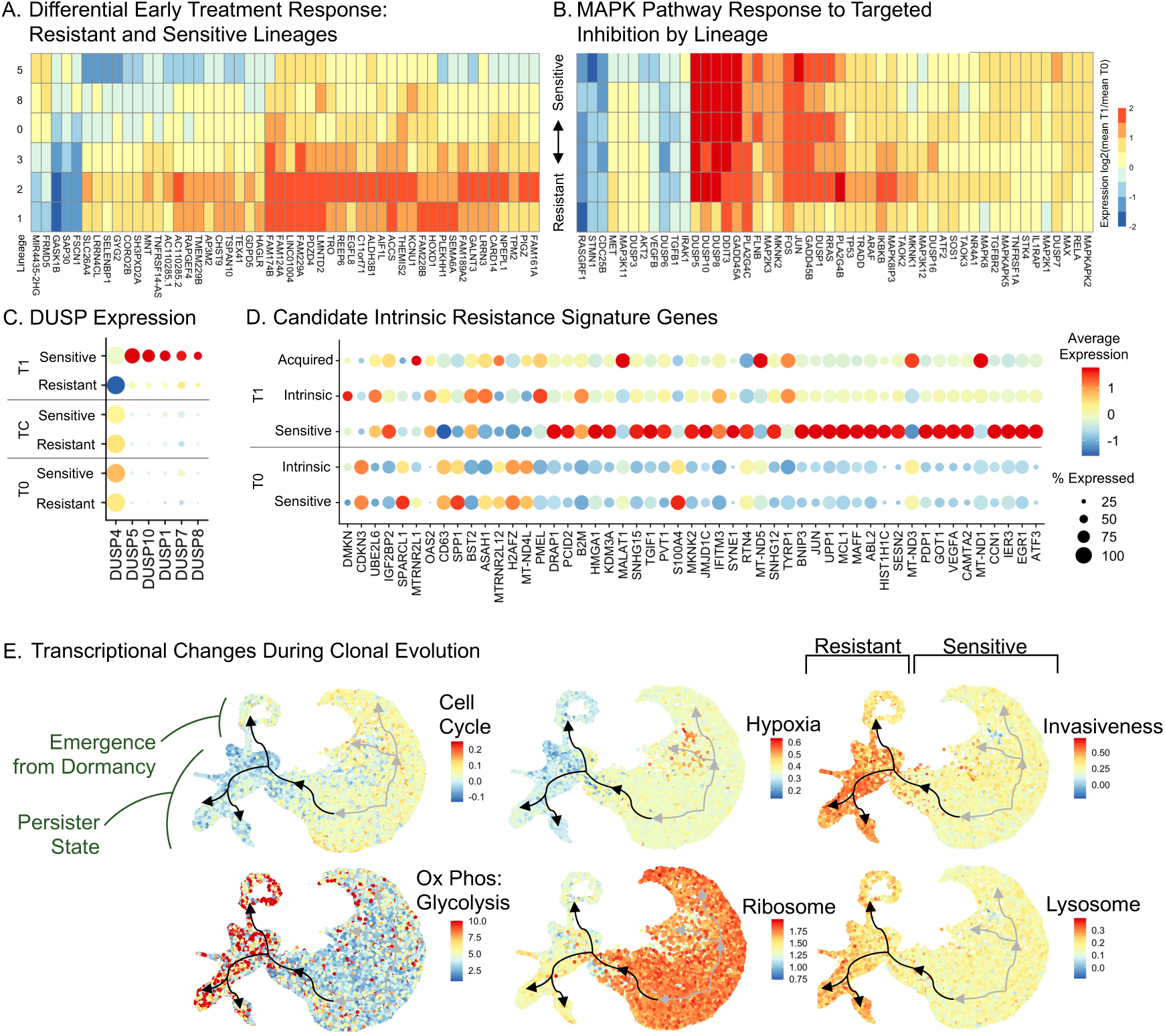
Temporally resolved phylogeny tracing identifies intrinsic and acquired resistance signatures with potential therapeutic targets. A. Top differentially expressed genes between time point 1 and time point 0 in the resistant lineages (1,2) but not in the sensitive lineages (0,5,8) (Bonferroni corrected p-values < .0001).[JC1] Lineage (3) is the ancestral lineage. Data are for WM4007. B. Top differentially expressed MAPK-pathway genes by lineage, T0 vs. T1. C. Treatment induces upregulation of DUSP genes in sensitive but not in resistant lineages.[JC2] D. Differentially expressed genes between intrinsically resistant and all other lineages. Expression levels show differential response to treatment between sensitive, intrinsic-, and acquired-resistance lineages. E. Plots of cell cycle, hypoxia, invasiveness, ox phos : glycolysis, ribosome, and lysosome on the CNV UMAP for WM4007. Transcriptional changes during emergence from dormancy show decreased invasiveness and continued reliance on oxidative respiration. Abbreviations: Uniform Manifold Approximation and Projection (UMAP), Copy Number Variant (CNV)

This phylogenetic reconstruction also divides the UMAP into treatment sensitive vs. resistant spots (Figure 5d, right-middle panel) with CNV lineage 3 representing the common ancestor. The spatial locations of sensitive and resistant spots at T0 and T1 are noteworthy (Supplementary Figure 6). The resistant lineages lie closer to the tumor periphery in both T0 and T1, and by T1 these lineages appear to be outcompeting the sensitive ones. The untreated control sample demonstrates growth of both sensitive and resistant lineages, but in the absence of treatment the sensitive lineages are able to compete successfully. Strikingly, in the untreated samples CNV lineage 3, the root of the phylogeny, often occurs at the spatial boundary between the sensitive and resistant lineages. These patterns support a spatially determined mechanism of resistance, that the mechanism is acquired, and that the acquisition is dependent upon local conditions and interactions at the tumor periphery. Taken together, the juxtaposition of the RNA-derived clusters with CNV-lineages shows that phylogenetic evolution is distinct from transcriptional state change. Induction toward the persistent state occurs globally, while emergence back into a proliferating state can only occur in resistant lineages, as the others are extinguished by treatment.

### Early response to treatment differs in sensitive and resistant lineages and identifies DUSP genes associated with resistance

The ability to distinguish sensitive from resistant lineages can clarify the transcriptional profiles that accompany tumor survival and regrowth. Comparisons of before-vs. after-treatment samples reveal expression changes that are lineage-specific. Genes showing significantly altered expression in T1 vs. T0 in the resistant but not the sensitive lineages are shown in Figure 6a, expressed as log2(T1/T0) (top 50 genes, Bonferroni adjusted p<.0001 for all). Cellular respiration and ribosome biogenesis are significantly enriched GO terms among these genes (adjusted p<.0001 for both). There are 823 genes with an opposite response to treatment in the sensitive vs. the resistant lineages. Among these genes, the ribosome, cell cycle, and oxidative phosphorylation KEGG pathways demonstrate highly significant enrichment (adjusted p<.0001 for all)(Supplementary Table 4). Comparison of all MAPK-pathway genes before and immediately after the initiation of treatment (T0 vs. T1) also identifies a differentially expressed subset with distinct responses in sensitive vs. resistant lineages (Figure 6b, Supplementary Table 5). These include several dual specific phosphatases (DUSPs), which are known to cause downstream suppression of ERK and JNK in the cell nucleus leading to decreased proliferation and differentiation, and our findings support prior studies on their potential importance in resistance mechanisms.^20–22^ DUSP proteins have also been noted to play a role in cellular plasticity and melanoma drug resistance.^23^ We find significant upregulation of several DUSP family genes (DUSP 1, 5, 7, 8, 10) upon initiation of treatment in the sensitive but not the resistant lineages, whereas DUSP4 expression is strongly suppressed in the resistant but not the sensitive lineages (Figure 6b/c, Supplementary Figure 7). These DUSP effects are not seen in comparisons of the untreated-early (T0) vs. untreated-late (TC) samples.

### Temporally resolved phylogeny tracing distinguishes intrinsic and acquired resistance signatures

A small population of lineage 1 in model WM4007 exists prior to treatment, peaking in size at T1 while also undergoing a transcriptional state change. Although lineage 4 derives from lineage 1 (Figure 5b,c), it is noteworthy that lineage 1 itself survives through time point 4, as this implies a level of intrinsic resistance. At T1, comparison of lineage 1 to all others identifies 51 differentially expressed genes (adjusted p<.05 for all), with enrichment in the oxidative phosphorylation pathway (p<.05), again demonstrating metabolic symbiosis with glycolytic respiration as a mechanism of melanoma plasticity (Figure 6d, Supplementary Table 6). This behavior comports with the metabolic shift that has been explored as a possible target using novel specific inhibitors^24^ and that has supported medications such as metformin for the melanoma armamentarium.^25,26^ Additional, non-oxidative phosphorylation related genes show divergent treatment response, such as Reticulon-4 (RTN4) which is significantly upregulated in the sensitive lineages and downregulated in the resistant lineages at T1. Decreased expression has been found to increase susceptibility to chemotherapy both *in vitro* and *in vivo*. These findings suggest that knowledge of the temporal dynamics of cancer cell state change through the treatment course could identify therapeutic windows where agents without a traditional role in melanoma treatment may gain efficacy.^27^

Although a small population of lineage 1 survives via intrinsic resistance, the majority appears to evolve into lineage 4, which can be analyzed to distinguish acquired resistance mechanisms. Comparison of lineage 4 to the other lineages identifies 1166 DEGs; the subset of these DEGs in lineage 4 that are not in lineage 1 provide transcriptional signatures of acquired resistance (Supplementary Table 7). The FoxO (p<.01) and p53-pathways (p<.0001) are significantly associated with this gene set (Supplementary Figure 7). An example gene from these pathways of interest as a potential therapeutic target is CDK2, which is down-regulated in the sensitive lineages and upregulated in the resistant. Indeed, evidence exists suggesting efficacy of specific CDK2 targeting in melanomas resistant to BRAF inhibition.^28^

We observed different patterns of evolution related to prior treatment history. The model described above (WM4007) derives from a treatment-naive tumor, whereas model WM4237 comes from a patient who received two rounds of immunotherapy prior to tumor procurement. The phylogenetic tracing of WM4237 indicates pre-existing resistance mechanisms common to most lineages (Supplementary Figure 4). All lineages existing prior to treatment survive to time point 4 and undergo global transition to the persister transcriptional state, albeit at different tempos. Unlike WM4007, we observe no lineage extinction due to treatment in WM4237. Some resistant lineages (4, 5, 8, 10, and 11) emerge under treatment but are negligible in the control sample. Meanwhile, resistant lineages 0, 1, 2, 3, and 6 already dominate the pre-treatment and the control samples, and persist in spite of treatment even at T4, though at attenuated proportions. In contrast, resistant lineage 7 comprises a small population pre-treatment, but shows significant expansion both with and without treatment. Finally, resistant lineage 9 appears to have a therapy-dependent fitness advantage, expanding under the selective pressure of treatment, but not in the control sample.

Lineage analysis in WM4237 reveals commonalities in resistance mechanisms with model WM4007. Notably, several lineages in WM4237 (predominantly lineage 10, which is closely related to 1) begin to expand in T4 while simultaneously returning to pre-treatment expression states (RNA clusters 0, 1, 2, 5, 6) (Supplementary Figure 4). Although at T4 these cells return to a transcriptional state similar enough to cluster with pre-treatment cells, there are 1,673 genes that are significantly differentially expressed between T4 and T0, including 40 of the 301 genes in the MAPK pathway (Supplementary Table 8). Among the differentially expressed MAPK genes are members of the DUSP family (DUSP 4, 5, 6, 10, 15), which show decreased expression early in treatment followed by upregulation in the late-arising, proliferating lineage. A subset of these genes (DUSP 4, 5, 10) are also differentially expressed in the resistant versus sensitive lineages in model WM4007 (Figure 6c).

### Transcriptional changes during emergence from dormancy reflect decreased invasiveness and continued reliance on oxidative respiration

Comparison of populations in the persister state (e.g. WM4007 T3 lineage 4) with the definitively resistant populations (e.g. WM4007 T4 lineage 7) reveals the transcriptional shift accompanying the return to a proliferative state (Figure 5b). Comparison of these lineages identified 2,924 DEGs (Supplementary Table 9). Pathway analysis on this gene set confirms that emergence from the persister state is accompanied by an increase in the cell cycle (p<.00001), oxidative phosphorylation (p<.00001), and p53 signaling KEGG pathways (p=.002) (Figure 6e). This finding suggests that targeting mechanisms of oxidative respiration may indeed complement traditional targeted therapy, though it raises the question of whether such targeting would kill the respiring population or epigenetically drive it back to glycolysis. As noted above, invasive capacity, as reflected in the gene set score derived from Hoek *et al*,^29^ was increased in the dormant persister state. However, invasive capacity was markedly attenuated during the transition to the proliferating resistant lineage. This provides *in vivo* evidence to support prior *in vitro* studies postulating a mechanism where cells evade therapy in the slower cycling but more invasive state, but can transition back to a proliferative state at the cost of invasive capacity.^29^

Other pathways also show dramatic changes across the evolutionary bottleneck. Ribosome subunit expression is high in the early timepoints of WM4007, decreases in the persister state, and reaches yet lower levels in the resistant, proliferating lineage (p<.00001) (Figure 6e). Given that ribosome biogenesis allows for the increased protein synthesis capacity required of proliferating cancer cells, ribosomal subunits have been suggested as potential therapeutic targets in melanoma and other cancer types.^30^ Similarly, cancer cells that are proliferating rely heavily on efficient lysosomal function, and this has been explored as a potential therapeutic target.^31^ Converse to ribosomal expression, we note that lysosomal expression increases in the persister state and remains elevated in the resistant, proliferating lineage (p<.00001). These results suggest another potentially synergistic pair of therapeutic targets, highlighting the critical importance of timing specific agents such that the tumor is in a sensitive evolutionary stage and transcriptional state.

### Deep learning imaging features identify shifting transcriptional states and lineage evolution

Given the complexity of spatial expression profiling, we also investigated whether major transcriptional states can be detected by the technically simpler approach of analyzing H&E-stained images. These images are extracted from the same tumor sections used for the spatial transcriptomics (ST H&E). H&E-stained images are processed through deep learning pipelines, producing numeric imaging features spatially aligned with the transcriptomic data. Figures 7a,c show the UMAP projection of these imaging features, demonstrating a progression from the left to the right side of the projection during treatment, with a subpopulation shifting back toward the left in T4. The TC control sample remains at the left with the T0 sample. Imaging features segregate disease states– from pre-treatment to MRD to the emergence of the proliferating, resistant state. We also observe similarity between technical replicates, which were derived from H&E-stained sections taken at alternative locations within the block and processed at different histology facilities (see Methods). This implies that H&E quantifications are consistent at regions proximal within the tumor and robust to variations in slide preparation and image scanning hardware (Supplementary Figure 8).

**Figure 7.**
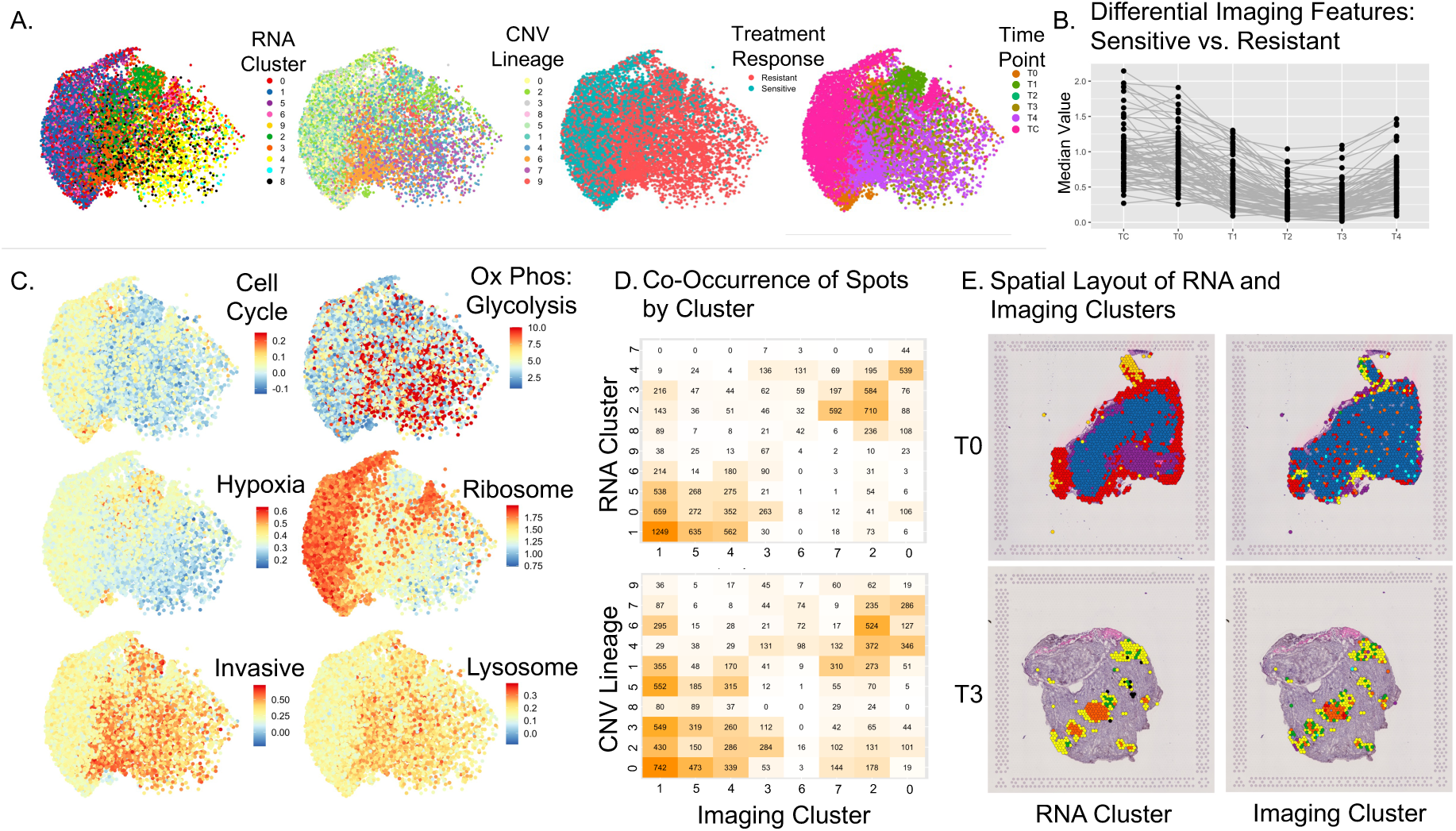
Imaging features identify shifting transcriptional states and evolutionary lineage (WM4007) A. UMAP projection of imaging features for each ST spot, colored by RNA cluster, CNV lineage, resistance potential, and time point (left to right). These show a progression from the left to the right side of the projection as treatment progresses, except for a subpopulation shifting back toward the left at time point 4. B. 77 differential imaging features that segregate resistant from sensitive tumor spots, plotted by treatment time on the x-axis. These show dampening of the features’ signals across treatment time with a late uptick as cell proliferation begins anew. C. UMAP projection of imaging features for each ST spot, colored by expression scores for notable gene sets. These gene sets are well-localized within the image feature UMAP, analogous to their localization within the CNV UMAP (Fig 5e). D. Co-occurrence matrix of imaging-derived clusters vs. (top) rna-derived clusters and (bottom) cnv-derived lineages. E. Spatial layout of RNA and imaging-defined clusters at timepoints T0 and T3 in WM4007. Abbreviations: Uniform Manifold Approximation and Projection (UMAP), Copy Number Variant (CNV)

Imaging features also segregate with the CNV-defined treatment-sensitive versus - resistant lineages. The treatment sensitive spots are extinguished relatively early and can be seen to overlie the untreated and early treatment time points on the left side of the imaging UMAP (Figure 7a, right-middle panel, Supplementary Figure 8). The resistant spots span both sides of the UMAP, demonstrating that the histopathological appearance of resistant clones changes during the course of treatment, migrating to a distinct appearance during the persister state and then back toward the initial pre-treatment morphology. However, as the resistant spots return toward this initial morphology, they do occupy distinct regions on the left side of the UMAP. This indicates that while they shift away from their persister appearance, they maintain an imaging signature distinct from their pretreatment counterparts. Of interest, individual deep learning features can be identified that drastically vary upon initiation of treatment. Differential image feature analysis identifies 77 candidate features that segregate resistant versus sensitive tumor spots. These exhibit a dampening of the imaging feature amplitudes across treatment time, though with an uptick as cell proliferation begins anew (Figure 7b). Superposition of gene set scores on the imaging-derived UMAP shows good separation of spots on the basis of the same pathways noted above to vary significantly by CNV lineage (Figure 7c).

Comparison of spot clusters defined by gene expression to clusters defined by imaging features demonstrates strong correlation between the two modalities (Figure 7d). Imaging clusters 1, 4, and 5 show particularly clear association with the RNA clusters dominating the untreated tumor state. In contrast, imaging clusters 0, 2, and 7 are aligned with RNA clusters dominant in the minimal residual disease state. The correlation between CNV Lineage and imaging clusters is analogously apparent albeit not as strong, indicating that epigenetic state is more closely associated with phenotypic appearance than is phylogenetic lineage. These associations demonstrate the ability of imaging features to distinguish genomics-derived groupings. Visualization of the imaging-defined clusters alongside RNA clusters within tissues shows that deep learning H&E analysis can distinguish the central-to-peripheral expression gradient in the pre-treatment sample. The correlations between H&E and expression state are even more pronounced with respect to the critical persister state as seen in T3 (Figure 7e).

## Discussion

Tumor clonal populations are known to enter a dormant, persister state upon exposure to treatment with the eventual emergence of proliferating, resistant lineages.^8,32–34^ However, due to difficulties of longitudinal sampling of tumors during the treatment course, much is unknown about the mechanisms of entry into and emergence from the persister state, the prior presence or development of resistance capability, and the interplay between heterogeneous clonal lineages. In this study, we track shifting expression signatures through the treatment course while simultaneously tracing the phylogenetic evolution of clones with varying capacity for therapeutic resistance. We find that imposing the selective pressure of targeted treatment upon the inherently heterogeneous clonal populations comprising BRAF mutant melanoma sets in motion a programmatic response with reproducible spatial and temporal patterns.

Previous work has described the phenotypic plasticity of the persister state, which has been characterized as slow-cycling and involving reversible upregulation of histone demethylases.^4,6,7^ Additionally, the persister state has demonstrated shifts in the mechanism of cellular respiration and emphasis on the clearing of free radicals.^19,34–39^ However, many of these findings result from experiments involving cell culture, with artificial induction of a persister-like state and a lack of spatial and temporal resolution. Our PDX model parallels the treatment-driven evolution occurring in the clinical setting while maintaining structural relationships within the tissue and provides validation for the previously identified cell state shifts occurring during entry into the persister state. Our longitudinal treatment design demonstrates that dense time point sampling, particularly in the early treatment stages, is critical to capturing not just differences between the pre-treatment and MRD populations, but in elucidating the events driving transition into this persister state.

We further augment current understanding of the persister state by identifying trends in the spatial patterning among previously observed transcriptional shifts, finding interface populations of cells in a more stem-like state as well as recurrent central-to-peripheral gradients. The central-peripheral patterns suggest a tumor microenvironmental influence, where populations at the tumor-stromal boundary are subject to different signaling compared to the intra-tumoral regions, thereby driving alternative transcriptional changes. In addition to this possibility is the likelihood of increased ischemia and decreased exposure to the therapeutic agents in the central tumor regions, leading to differential cell state shifting at the tumor core, contributing to the plasticity allowing cells to enter relative dormancy. An experimental design using spatial but no temporal resolution provides only a snapshot of tumor behavior, markedly limiting the ability to capture the complex evolutionary dynamics driving treatment response. In contrast, using temporal but not spatial resolution could identify global shifts in expression and define clonal populations, but would lack insight into the spatially stereotyped nature of transcriptional change, be it in the clone-clone interfaces or central-to-peripheral gradients. Further understanding of the microenvironmental interactions occurring during treatment will be critical to elucidating the specific pathways driving transcriptional change, which in turn will help in the design of rational therapeutic strategies that stand a chance at longitudinal control of cancer progression. Such progress may depend on the application of experimental systems that provide clonal lineage and expression state information with both spatial and temporal resolution.

The concept of improving therapeutic efficacy through specific targeting of processes known to be associated with the persister state is attractive.^40–43^ However, without a more complete understanding of the dynamics of the expression state change and the evolutionary interplay occurring during treatment, such efforts are likely to fail at the hands of the highly adaptable tumor milieu. One repeated finding in the literature is that the ability to enter the persister state is limited to a small subset of cells.^4,18^ A recent study applying chemotherapeutic agents to PDX models of colorectal cancer challenged this concept, reporting that, in fact, any cancer cell has the ability to undergo this state change.^19^ Our longitudinal treatment design with dense time point sampling and spatially resolved data adds to the evidence that persister ability is not restricted to a subset of cells and expands this finding to include response to targeted therapy in addition to chemotherapy. Furthermore, we capture a tumor in the midst of transitioning into the persister state and identify spatial patterns underlying this transcriptional change. By tracking evolutionary dynamics among clonal lineages entering this treatment bottleneck, we confirm that despite phylogenetic heterogeneity, the initiation of treatment does induce all tumor cell populations toward the persister expression state. We further delineate variable ability among the clonal lineages to survive the bottleneck within this state, confirm recent findings of multiple different pathways through the minimal residual disease state,^9^ and provide evidence that the re-emerging resistant clones were present prior to the initiation of treatment rather than arising de novo. We find that while resistance mechanisms are likely tumor specific and the genes involved are highly variable,^44^ there are MAPK-dependent and -independent pathways that support previous mechanistic findings. Specifically, the dual specific phosphatase (DUSP) family represents a promising potential target for intervention.^23,23^

Finally, the application of deep learning image analysis identifies imaging features that correlate with tumor state, treatment sensitivity, and invasive capacity. The addition of these digital pathology techniques to the experimental armamentarium brings the promise of increased understanding of the dynamic changes occurring during the treatment course, converting H&E-stained images generated from any biopsy specimen into an opportunity to derive spatially-resolved information regarding tumor state and evolutionary direction.

## Acknowledgments

This work was made possible by the contributions of the JAX Single Cell Biology Laboratory. We are also grateful for contributions from the Computational Sciences Core at the The Jackson Laboratory. Portions of this work were funded by support from Hartford Healthcare. Work performed by the Herlyn group is supported by the following grants: U54CA4224070, P01CA114046, P50CA261608, R01CA258113, and the Dr. Miriam and Sheldon Adelson Foundation. Work performed by the Chuang group was supported by R01 CA230031 and U24CA224067.

## Author Contributions

Conceptualization: J.C.R, M.H., J.H.C; Investigation: J.C.R., S.D., T.B.S., S.P., J.K., H.L.; Formal analysis: J.C.R., S.D., T.B.S., B.S.; Software: S.D.; Methodology: J.C.R., S.D., H.L., M.H., J.H.C; Resources: T.B.S, S.P., J.K., H.L., M.H.; Writing: J.C.R. and J.H.C. with input from all co-authors; Visualization: J.C.R.; Project Administration: J.C.R.; Funding acquisition J.C.R. and J.H.C.; Supervision: J.C.R., O.A., M.H., J.H.C

## Inclusion and diversity

We support inclusive, diverse, and equitable conduct of research.

## Supplemental information

Supplementary Tables (available at https://drive.google.com/drive/folders/1QH9noLfjGuKeA1yg8-ZuDjNj5xKKREfp?usp=drive_link)

Supplementary Table 1. Differentially expressed genes between pre-treatment (T0) and early-treatment (T1) that are significant in both models WM4007 and WM4237.

Supplementary Table 2. Individual genes and gene sets that exhibit significant correlation between expression level and distance from the tumor boundary.

Supplementary Table 3. Differentially expressed genes between the boundary cluster 7 (yellow) and both its internally and externally neighboring clusters, 4 (orange) and 3 (cyan)(WM4237).

Supplementary Table 4. Differentially expressed genes between time point 1 and time point 0 with significant corrected p-values in the resistant lineages (1, 2) but not in the sensitive lineages (0, 5, 8), in the sensitive but not resistant, and in both sensitive and resistant (WM4007).

Supplementary Table 5. Differentially expressed MAPK-pathway genes before and immediately after the initiation of treatment (T0 v T1)(WM4007).

Supplementary Table 6. Differentially expressed genes within T1, comparing lineage 1 to all others (WM4007).

Supplementary Table 7. Differentially expressed genes within T1, comparing lineage 4 to all others and excluding significant DEGs from the T1 lineage 1 to the rest comparison (WM4007).

Supplementary Table 8. Differentially expressed genes comparing T4 to T0 for RNA clusters 0, 1, 2, 5, 6 (WM4237).

Supplementary Table 9. Differentially expressed genes comparing T3 lineage 4 to T4 lineage 7 (WM4007).

Supplementary Table 10. PDX mice identification, time of tumor tissue harvesting after the beginning of treatment, and treatment regimen summary.

## Methods

### 1. Experimental Design

The two WISTAR PDX models, immunotherapy-treated WM4237-1 and treatment-naive WM4007, are BRAF V600E mutant metastatic melanomas of a subcutaneously implanted solid tumor on NSG mice. Twenty mice per model were used in the longitudinal experiment. When the tumors reached 300 mm^3^, measured biweekly, the continuous dose treatment of mice via chow with either a dual BRAF/MEK inhibitor or vehicle was initiated. The mice were either untreated, treated with phosphate-buffered saline with 5% DMSO, or treated with dual BRAF/MEK inhibitor dabrafenib (150mg/kg)/trametinib (1.5mg/kg) via chow (∼5 g/day).

One untreated mouse per model was sacrificed at time point T0, day 0. One vehicle-treated mouse per model was sacrificed at time point TC, days 42 and 38, WM4237-1 and WM4007, respectively. The tumors of therapy-treated sacrificed mice, one tumor tissue block per mouse per model per time point, were harvested at time points T1 (day 14), T2 (days 42 and 44), T3 (days 70 and 73), and T4 (days 94 and 133). We analyzed two near adjacent tissue sections from each tumor tissue block for each model and time point using H&E staining profiling and 10x Visium Spatial Gene Expression.

### 2. Tissue Sectioning and H&E staining

Cryotome-cut OCT-embedded fresh frozen tissue was used to generate tissue sections on microscopy slides. Two sections were prepared per tissue block, then covered with mounting media and coverslip. The glass slides were scanned on Aperio Leica Biosystems GT450 scanner at 40x magnification and a resolution of 0.263748 microns per pixel (mpp), compressed with JPEG/YCC Q=91. Regions of interest containing any tissue were extracted from whole slide images and converted to uncompressed RGB TIFF images.

### 3. Visium Profiling

The Jackson Laboratory Scientific Services, Single Cell Biology Laboratory, generated the assay. OCT-embedded fresh frozen tissue was used. Five 10µm sections from each block were used for total RNA extraction (Qiagen), which was used for RNA integrity determination by RINe score using an Agilent Tapestation and High Sensitivity RNA ScreenTape. Blocks with RINe greater than 7 were optimized for ideal permeabilization time following the vendor protocol (10x Genomics, CG000238). Sections from four tissue blocks were placed on Visium Gene Expression slide for H&E staining and brightfield imaging via NanoZoomer SQ (Hamamatsu), block-specific tissue permeabilization, mRNA capture, and subsequent library generation per the manufacturer’s protocol (10x Genomics, CG000239). Whole slide images of the Visium slides were split into uncompressed full-resolution RGB TIFF images of capture areas via OMERO Python API.

### 4. Visium Raw Sequencing and H&E Image Data Processing

Illumina base call files for all libraries were converted by The Jackson Laboratory Scientific Services, Single Cell Biology Laboratory, to FASTQ files using bcl2fastq v2.20.0.422 (Illumina).

### 4.1. Data Pre-processing with the STQ pipeline

FASTQ files were split into the mouse and human counterparts with “xenome classify” of Xenome 1.0.1.^45^ Xenome classification indices were prepared with “xenome index” of Xenome 1.0.1, Human GenBank assembly GCA_009914755.4 of T2T-CHM13v2.0 genome reference, and non-obese diabetic (NOD) mouse genome assembly. Human sequencing reads were aligned to GRCh38-2020-A (GENCODE v32/Ensembl 98) genome with “spaceranger count” of Space Ranger 1.3.1, mouse sequencing reads were aligned to mm10-2020-A (GENCODE vM23/Ensembl 98) genome. The two reference genomes were downloaded from the 10x Genomics data portal. Human and mouse gene count matrices were merged into one matrix. Image and ST spots grid alignment was done with Space Ranger 1.3.1.^46^ The details of pre-processing steps implementation are available in the STQ pipeline.^47^

### 4.2. Quality Control and Normalization

Spatial Transcriptomics pipeline (https://nf-co.re/spatialtranscriptomics, fork “spatialtranscriptomics” at https://github.com/sdomanskyi/) was used to QC, normalize, and inspect the Visium data from each sample. We applied filtering options in the pipeline to keep all expressed genes and ST spots with at least 500 UMI counts and at least 250 genes expressed. For normalization, we applied variance stabilizing transformation (VST) implemented in the Seurat^48–51^ and corrected the spot by gene UMI count matrix with the transformation for each of the samples.

Before scaling each sample’s spots library sizes, we computed the sum of UMI counts of human genes per spot. Then we computed the median library size, 𝑚_!_, per sample using the spots with at least 80% human reads. For each sample, we scaled the distribution of spots’ library sizes by multiplying UMI counts by 20,000/𝑚_!_. We added 1 to the scaled UMI counts and log-transformed the values, and concatenated samples of each model with the union of the genes expressed in the samples of each model.

We calculated the Pearson correlation coefficient of average UMI counts values per gene across samples in two ways: by comparing the technical replicates (same time point) and biological replicates (different time points), see Supplementary Figure X.

#### 4.2.3. Filtering of mouse stroma spots

To determine ST spots containing cells of mouse origin (i.e., stroma), we subset the normalized data to mouse gene UMI counts and do dimensionality reduction as follows. We used the package scanpy^52^ to determine the top 2500 highly variable imaging genes (HVG) with a batch key “sample” and perform principal component analysis (PCA) on centered HVG to obtain 100 principal components (PCs). We corrected the PCs with harmony^53^ with the default settings of the package harmonypy^54^. We built a k-nearest neighbors (kNN) graph from the harmonized PCs and clustered spots with a high resolution of (1.0 for WM40237-1 and 0.75 for WM4007) by using data from all samples in each model. Clusters containing primarily spots with high mouse-to-human counts ratio and clusters covering skeletal muscle (applicable to samples of model WM4007) were marked as mouse stroma; see Supplementary Figure X.

#### 4.2.4. Dimensionality reduction, batch effect correction, clustering

We removed any ST spots marked as mouse stroma or bubbles from detected imaging artifacts, Section “Imaging features analysis.” We determined the top 2500 highly variable imaging genes (HVG) with a batch key “sample” and performed principal component analysis (PCA) on centered HVG to obtain 50 principal components (PCs). We corrected the PCs with harmony^53^ with default settings, except parameter lambda was set to 0.5 of the package harmonypy^54^. We built a k-nearest neighbors (kNN) graph from the harmonized PCs and clustered spots with a high resolution of (0.75 for WM40237-1 and 0.55 for WM4007) by using data from all samples in each model.

### 4.3. Differential gene expression analysis, pathway over-representation (ORA) analysis

To find differentially expressed genes across identifiers of interest, we used the Seurat function “FindAllMarkers” with the Wilcoxon Rank Sum test, a minimum fraction of spots expressing a gene 0.10, log-fold change threshold 0.15, and subsampling spots to a maximum of 500 per label. We retained differentially expressed genes with p-values corrected for FDR by the Bonferroni method under 0.05. The lists of differentially expressed genes were input to Metascape^55^ web application, an Overrepresentation Analysis (ORA) tool built on an extensive collection of ontology databases. We ran Metascape in “Multiple Gene Lists” mode and the default analysis option, “Express Analysis.” Pathways of interest, determined from ORA analysis and literature, were downloaded from Molecular Signatures Database (MSigDB).^56^ We use the gene sets with GSEApy, a Python implementation of GSEA (Gene Set Enrichment Analysis) and wrapper for Enrichr,^57^ to perform GSEA and single sample GSEA (ssGSEA).

### 4.4. RNA Velocity Analysis

RNA splicing quantification was done with Velocyto^10^ integrated into the STQ pipeline, which generated LOOM data files with reads classified as spliced, unspliced, or ambiguous. We filtered out ST spots with up to 25% of mouse reads and discarded the ambiguously spliced reads. We used scVelo^58^ to generate the velocity graph and velocity stream embeddings for visualization. We used scVelo to rank velocity genes and analyze individual genes’ velocity and velocity confidence.

To evaluate if there were divergence regions on the RNA velocity embedding maps, we developed an approximate calculation of RNA velocity (i.e., a vector field 𝐹) divergence, see Supplementary Figure X. Divergence of the vector field 𝐹 within a region relates to the two-dimensional flux of 𝐹 through the boundary of the region, 𝐶, a circle of radius 𝑅:

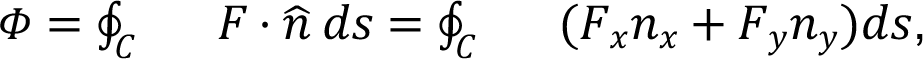

which we approximate with:

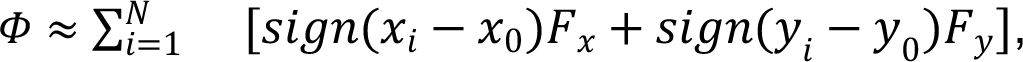

and use all values of 𝐹 within 𝐶 to estimate 𝛷, since the map of 𝐹 is not defined in every point of the velocity embedding. The flux is then projected on ST spots for spatial layout visualization and downstream analysis. The positive flux is indicating “sources” in divergence of the RNA velocity, or regions where RNA velocity is emerging from, while the negative flux indicates “sinks” of the RNA velocity.

### 4.5. CNV lineage inference

We used each model’s normalized and integrated gene expression data to estimate CNV frequencies with InferCNV^17^. We used spots of cluster 0 of timepoint T0 samples for WM4007 data and cluster 6 of samples of time point T0 for WM4237-1 as reference spots for inferCNV input. We calculated CNV burden as an average number of genes with a CNV event from the total pool of genes where significant CNV events were detected.

#### 4.5.1. Clustering, continuous ordering of CNV profiles of spots

Using package scanpy,^52^ we performed principal component analysis (PCA) on centered CNV profiles to obtain 100 PCs. We built a k-nearest neighbors (kNN) graph with metric “correlation” from the multiscale diffusion space with ten components, implemented in Palantir.^59^ We generated the UMAP representation of the kNN graph and performed Leiden clustering, with a resolution of 0.25, of the CNV profiles of spots from all samples in each model.

To infer continuous ordering of the CNV profiles, we generated a principal curve with package scFates.^60^ We set the root of the curve at the node where the reference gene expression value or a reference gene score value is the largest. In the Results, we describe the selection of the reference genes and scores, including the “Hoek invasive score.” This reference set is used to determine the pseudotime in Figure 4. We converted the principal curve to a finely segmented representation, 1000 segments, and inferred the pseudotime (continuous) order with scFates.

Invasive score is used for the pseudotime analysis because the concept of pseudo-time requires choosing a root or an origin in the cells’ ordering, and it is known that the invasiveness of tumor cells is an important property that allows cancer cells to undergo EMT and metastasize. We observe that the invasiveness score has a large pseudo-time gradient in each treated sample, except for T4 cluster 7, Figure 4A, and the observable gradient in the untreated samples, making it a suitable quantity for pseudo-time root determination. Notably, the average invasiveness is highest in MRD and lowest in untreated samples, suggesting that the therapy aimed at tumor reduction must also try to reduce the invasiveness of the residual cells.

### 4.6. Imaging features analysis

Whole slide images were processed with the STQ pipeline.^47^ The H&E-stained images of non-ST slides were processed with the Macenko^61^ stain normalization option. Imaging features were extracted with Inception V3^62^ convolutional neural network model developed and trained by Google with ImageNet^63^ database. We used a pre-trained HoVer-Net^64^ convolutional neural network model pre-trained on the PanNuke^65^ for simultaneous segmentation and classification of nuclei. Nuclear morphometric features were calculated for each nucleus and assigned to ST spots or image tiles.

Using the package scanpy,^52^ we determined the top 250 highly variable imaging features (HVIF) and performed principal component analysis (PCA) on centered HVIF to obtain 30 principal components (PCs). We built a k-nearest neighbors (kNN) graph from the PCs and clustered spots with a resolution of 0.35 by using data from all samples in each model, either WM4237 or WM4007.

To identify ST spots or tiles with imaging artifacts for each model’s H&E-stained tissue images, we performed PCA on all centered imaging features to obtain 30 PCs. We build a k-nearest neighbors (kNN) graph from the PCs and cluster spots with a resolution of 1.0, obtaining more than 10 clusters. The clusters containing air bubbles, tissue folds, excessive tissue tears, or loose stromal tissue were marked as imaging artifacts.

## 5. Whole Slide Images Post-processing

The whole slide images derived from either H&E-staining of tissue sections on microscopy slides or 10x Visium Spatial Gene Expression Slides were converted from uncompressed TIFF to pyramidal OME-TIFF, internally tiled with 1024 by 1024 pixels by using Open Microscopy Environment Bio-Formats^66^ tool bfconvert version 6.14.0. The executed command is “bfconvert -noflat -pyramid-resolutions [levels] -pyramid-scale 2 -overwrite” with 3-5 levels depending on the slide dimensions. The OME-TIFF files were deposited at images.jax.org, project ID 1451, with the help of JAX Research IT.

## Data and Code Availability

The analysis codebase and scripts are hosted at The Jackson Laboratory GitHub: https://github.com/TheJacksonLaboratory/PDX-melanoma-integrated-analysis. Raw and processed 10x Visium Spatial Gene Expression data are deposited at Gene Expression Omnibus (GEO) under accession GSE245582, project PRJNA1029019. The full-resolution images can be viewed or downloaded at https://images.jax.org/webclient/?show=project-1451. Detailed processed data is available on Zenodo (https://zenodo.org/records/10287542?token=eyJhbGciOiJIUzUxMiIsImlhdCI6MTcw MjMwNDEz[…]RCUzkVUo4Gp1G8EjFDOIEp8Zxug7XD00o7NvMSqEAol8Wl_DPvP Lx53IFmX6AZQ).

## Supplementary Figures

**Supplementary Figure 1.**
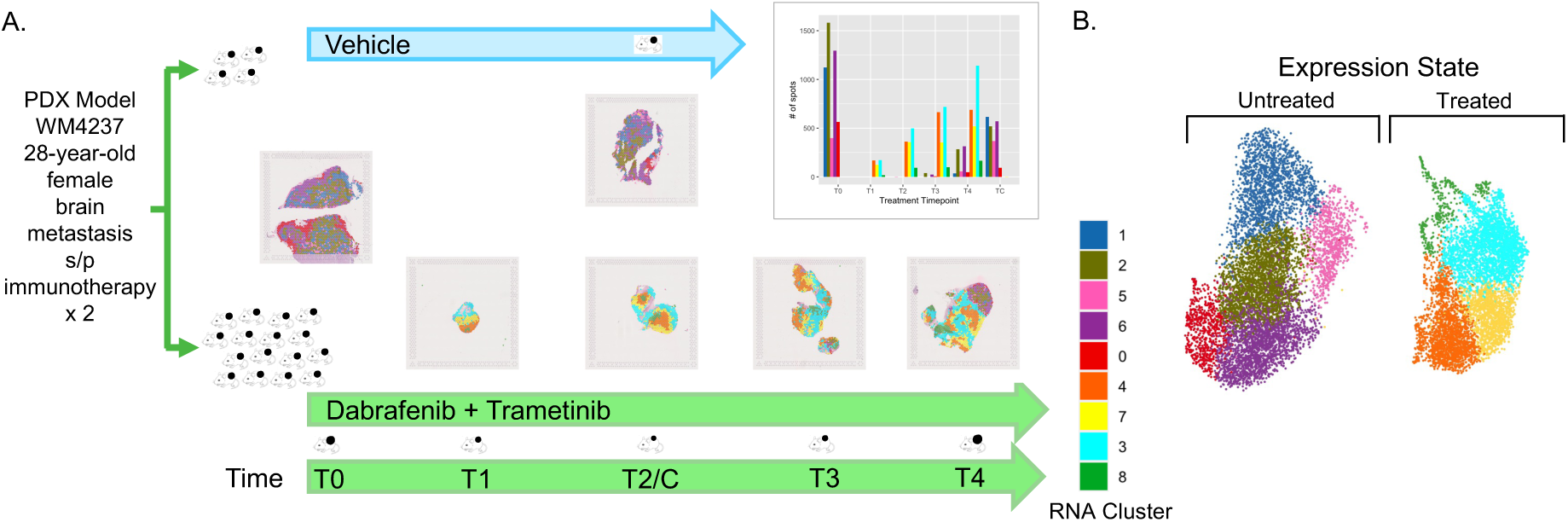
A. Summary of ST sections at each time point (WM4237), colored by RNA-derived clusters. Bar chart shows the proportion of RNA cluster per time point. B. UMAP projection of RNA clusters showing transition from a proliferating (left) into a more dormant expression state after treatment (right) Abbreviations: Spatial transcriptomics (ST)

**Supplementary Figure 2.**
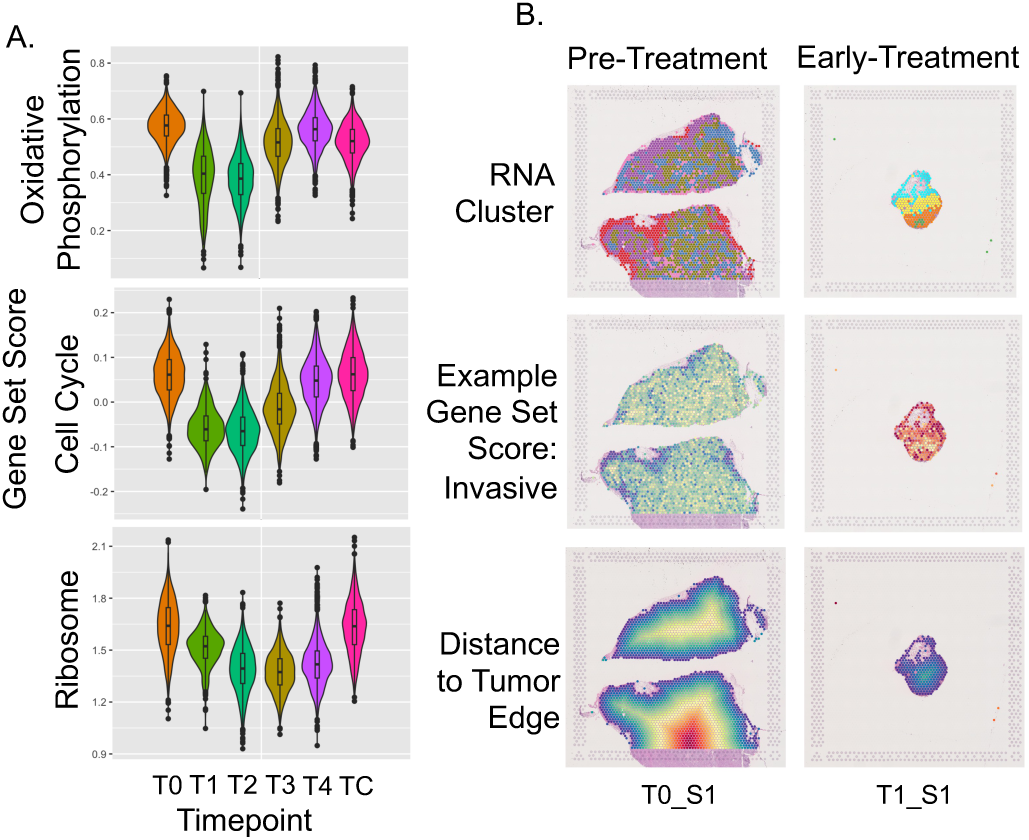
A. Pathway enrichment analysis of DEGs before and after the initiation of treatment (WM4237) demonstrates initial significant down-regulation of the cell cycle, oxidative phosphorylation, and ribosome pathways with return toward baseline in the late time points. B. Recurrent spatial patterns of RNA-derived clusters show specific expression states tending to occupy central tumor regions and others at the periphery or tumor-stromal boundary, more pronounced in the pre-treatment sample of model WM4237. The early treatment sample also shows a distinct spatial pattern.

**Supplementary Figure 3.**
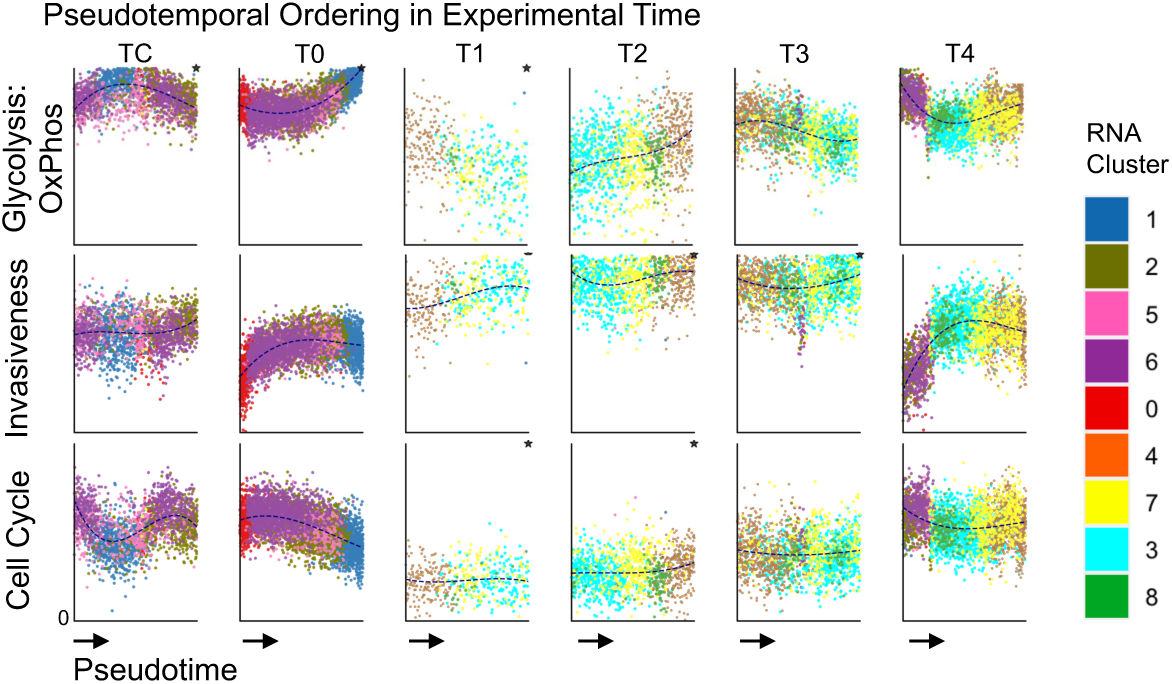
Metabolic plasticity and altered invasiveness-proliferation capacity mark transition into the persister state for WM4237. Pseudotemporal ordering reflecting the direction of transcriptional change, divided by treatment time point, with ST spots sorted vertically by gene set scores for glycolysis/oxidative phosphorylation, invasive capacity, and cell cycle (colors represent RNA-defined clusters). Treatment induces a shift from glycolysis toward oxidative phosphorylation as pseudotime progresses from cluster 4 to 7 to 3 (orange to yellow to cyan).

**Supplementary Figure 4.**
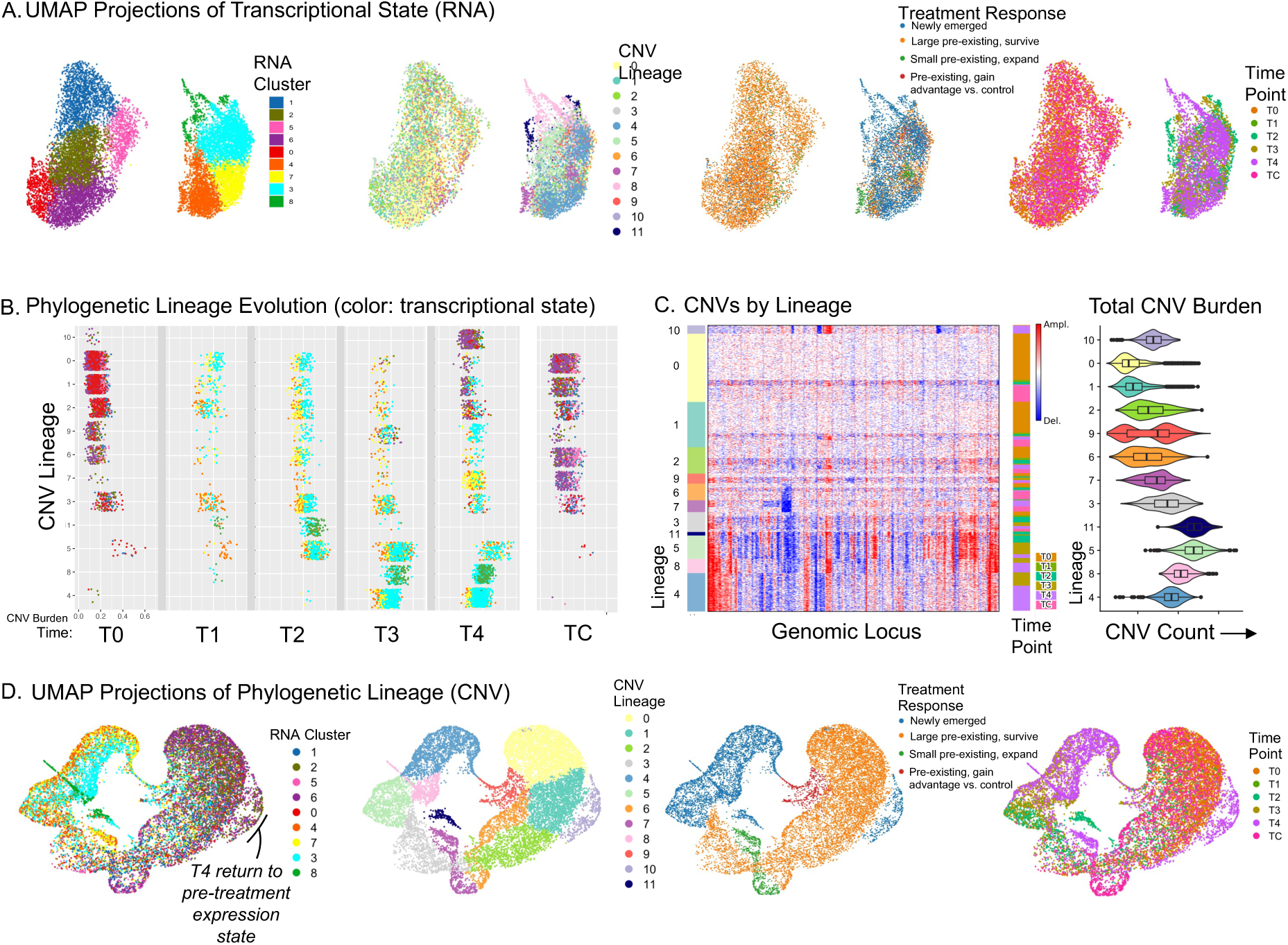
Phylogeny tracking reveals global transcriptional state change (WM4237). A. RNA-derived UMAP projection, colored by RNA cluster, CNV lineage, treatment response, time point (left to right). B. CNV lineage evolution reveals persistent-, emergent-, and expanded-lineages, as well as others that appear to have gained a selective advantage during treatment. Color shows RNA cluster, demonstrating transcriptional state change by lineage during treatment. C. (Left) Heatmap showing CNVs for each lineage, plotted for each spot and timepoint. (Right) Total CNV burden for each lineage across all spots, reflecting direction of evolution. D. CNV-derived UMAP projection, colored by RNA cluster, CNV lineage, resistance potential[JC5], and time point (left to right), demonstrating lineage evolution with late emergence of a resistant lineage closely related to those existing before treatment. Abbreviations: Uniform Manifold Approximation and Projection (UMAP), Copy Number Variant (CNV)

**Supplementary Figure 5.**
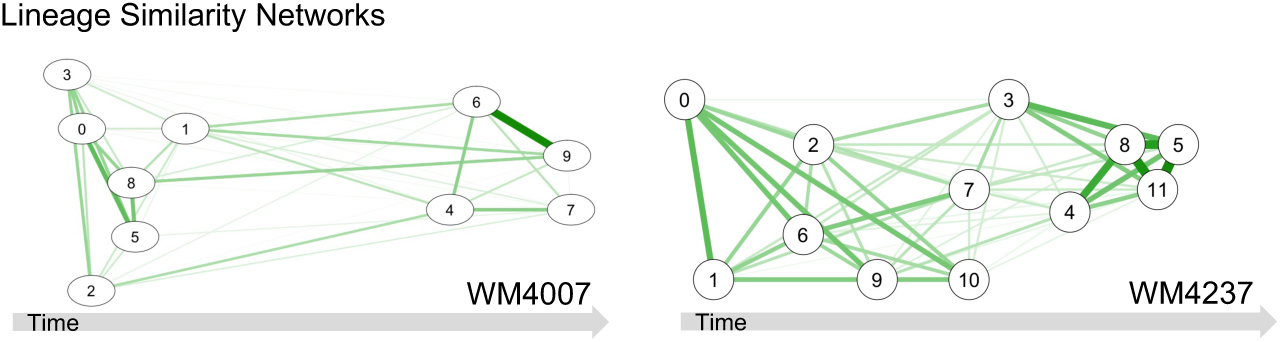
Lineage Similarity Networks for WM4007 and WM4237 show cosine distance between lineages based on CNV profile, with line weight representing degree of similarity.

**Supplementary Figure 6.**
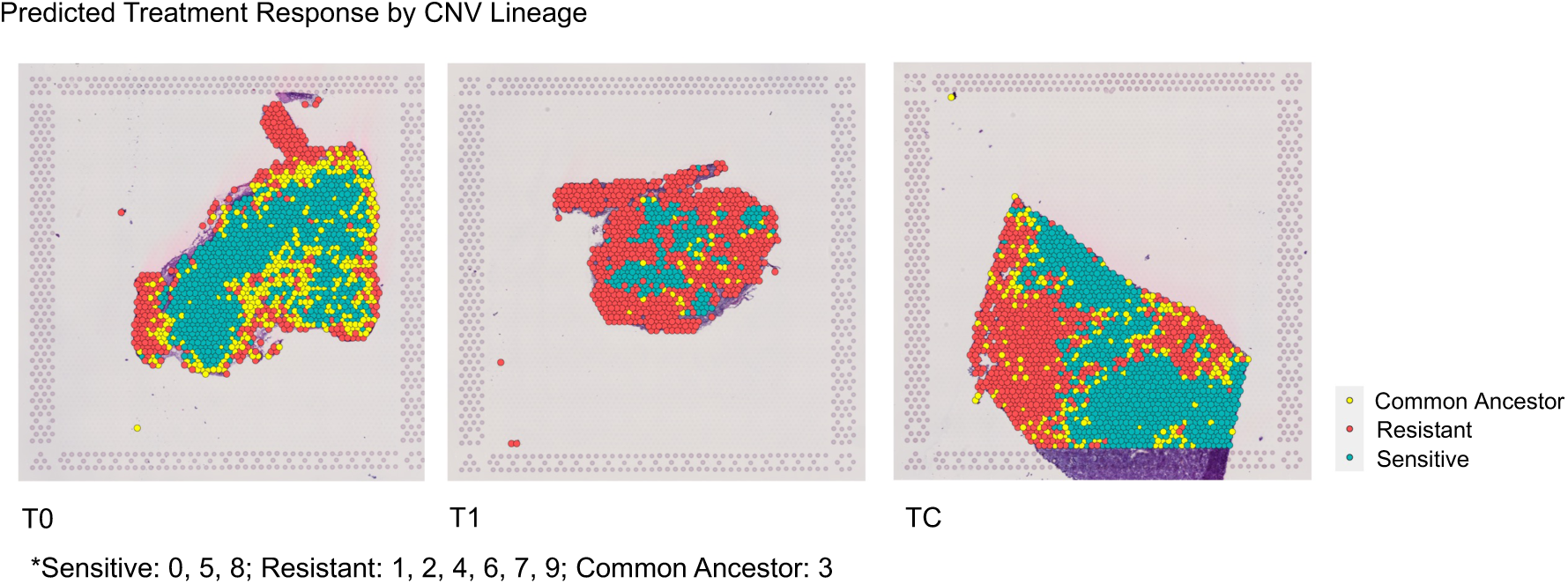
Spatial layout of WM4007 resistant vs sensitive lineages in time points 0, 1, and control demonstrate a boundary lineage that pre-dates the central and peripheral lineages. Sensitive, resistant and common ancestor lineages correspond to those labeled in Figure 5.

**Supplementary Figure 7.**
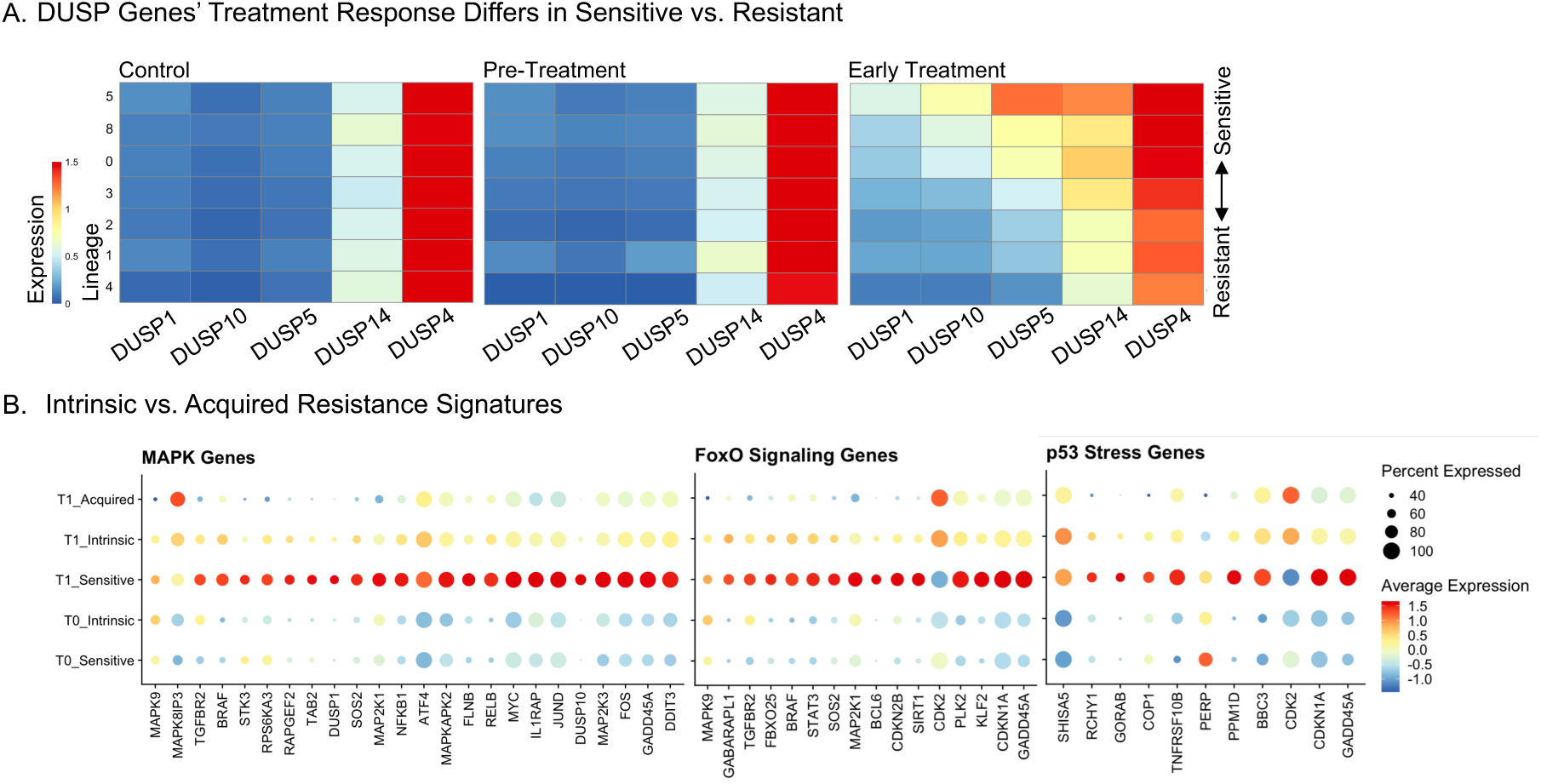
Differential treatment response in sensitive vs. resistant lineages. A. Differentially expressed DUSP genes from pre-treatment to time point 1 (WM4007) shows differential response in sensitive vs. resistant lineages, with down-regulation in the resistant lineages. B. Analysis on the set of differential expressed genes between intrinsic- and acquired-resistance lineages and sensitive lineages identifies significant enrichment of FoxO-, p53-, and MAPK-signaling pathways.

**Supplementary Figure 8.**
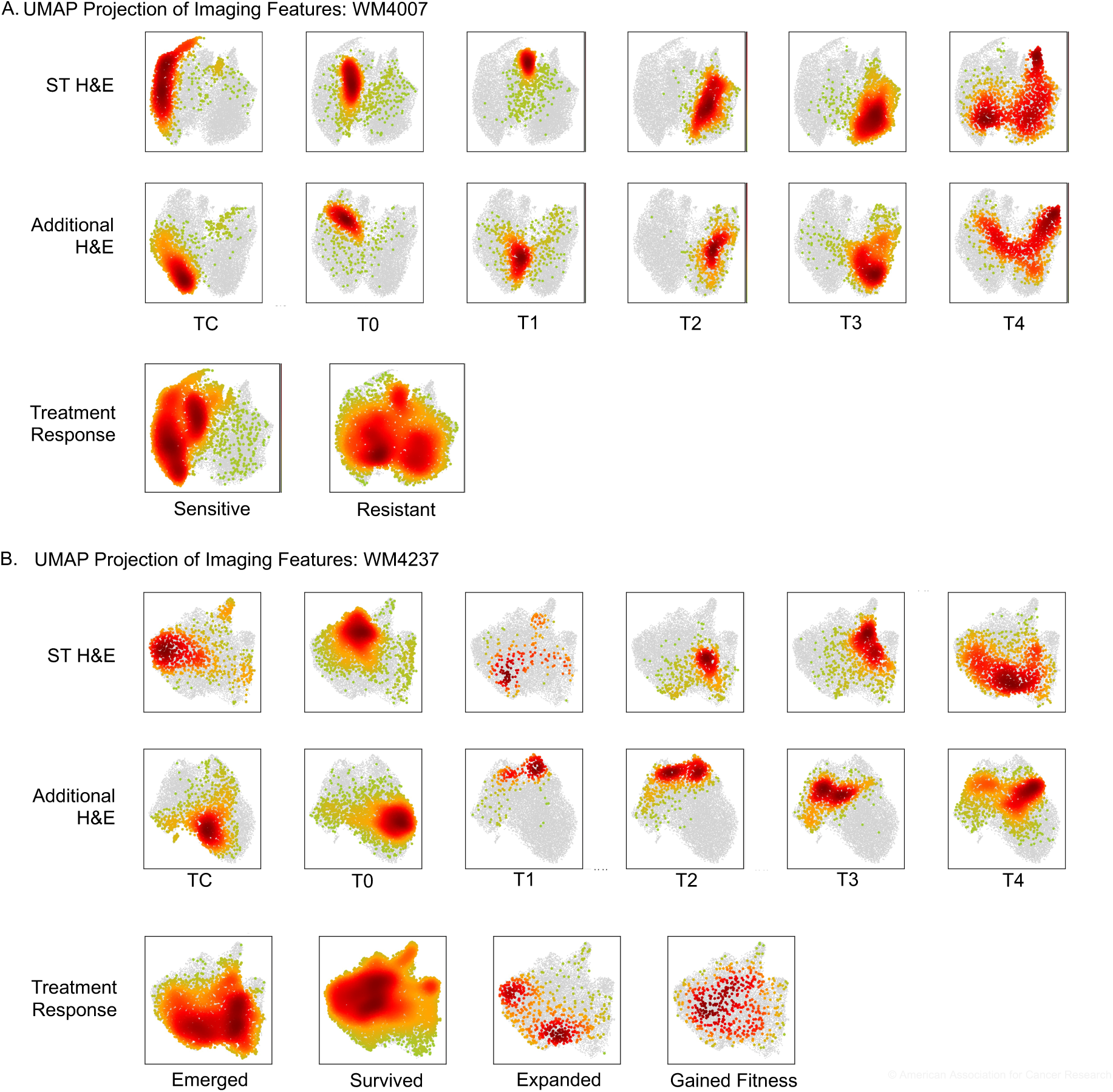
A. UMAP projection by imaging features (WM4007) from ST images and additional H&E-stained images taken from alternative location within the tumor block. The technical replicates each show progression of imaging appearance through treatment time. Resistant lineages separate from sensitive lineages in imaging features space. B. Same as above but for WM4237.

